# A Motion Correction Strategy for Multi-Contrast based 3D parametric imaging: Application to Inhomogeneous Magnetization Transfer (ihMT)

**DOI:** 10.1101/2020.09.11.292649

**Authors:** Lucas Soustelle, Julien Lamy, Arnaud Le Troter, Andreea Hertanu, Maxime Guye, Jean-Philippe Ranjeva, Gopal Varma, David C. Alsop, Jean Pelletier, Olivier Girard, Guillaume Duhamel

## Abstract

**Purpose:** To propose an efficient retrospective image-based method for motion correction of multi-contrast acquisitions with a low number of available images (MC-MoCo) and evaluate its use in 3D inhomogeneous Magnetization Transfer (ihMT) experiments in the human brain.

**Methods:** A framework for motion correction, including image pre-processing enhancement and rigid registration to an iteratively improved target image, was developed. The proposed method was compared to Motion Correction with FMRIB’s Linear Image Registration Tool (MCFLIRT) function in FSL over 13 subjects. Native (pre-correction) and residual (post-correction) motions were evaluated by means of markers positioned at well-defined anatomical regions over each image.

**Results:** Both motion correction strategies significantly reduced inter-image misalignment, and the MC-MoCo method yielded significantly better results than MCFLIRT.

**Conclusion:** MC-MoCo is a high-performance method for motion correction of multi-contrast volumes as in 3D ihMT imaging.

## Introduction

MRI techniques based on the combination of images with various contrasts for composite map generation are of great interest for structural (e.g. Diffusion Tensor Imaging, Magnetization Transfer) and functional (e.g. Arterial Spin Labeling) evaluations. Voxel misalignment across native images can however challenge the relevance of the parameters derived from these synthetic maps as they will translate into localized artefacts or loss of sharpness.

One of the main source of image misalignment comes from subject motion, which can occur during prolonged protocol durations or, in a more clinical context, during pediatric or pathological investigations (1–3). Compensation schemes to account for images misalignment must thus be considered. Although several retrospective dedicated motion correction functionalities exist and are publicly available (e.g. in FSL (4), ANTs (5) and SPM (6)), they are more suitable for acquisitions with a large number of images (e.g. serial repetitions in functional MRI) and little is available for parametric imaging techniques, which rely on the acquisition of a limited number of images with different contrasts.

Inhomogeneous Magnetization Transfer (ihMT) imaging is one such technique, which has proven to be of great interest for white matter assessment in the central nervous system (7–9). IhMT delivers a semiquantitative metric, the ihMT ratio (ihMTR), which is computed from a linear combination of serially acquired different MT-weighted images (MTw) normalized by a reference image acquired without MT saturation (MT_0_) (10,11).

The misalignment of images potentially occurring between image acquisitions during the images inter-acquisition essentially results in blurring and artefacts in the computed ihMTR map due to inconsistencies across combined voxels, which may prevent accurate analyses. This is particularly the case when image sharpness is essential, as for instance in the context of studying Multiple Sclerosis (MS) lesions, whose volumes scale from a few tens to thousands of cubic millimetres (7,9).

Both the small numbers of available images and different contrasts in an ihMT experiment limit the application of standard retrospective motion correction methods. In this work, we thus propose an efficient and robust motion correction framework adapted to the ihMT images features. The method is compared to the Motion Correction function FMRIB’s Linear Image Registration Tool (MCFLIRT) available in the FSL package (4). Both rigid-based approaches are evaluated before and after motion correction.

The proposed algorithm implementation and tools adapted for ihMT imaging are made freely available at: https://crmbm.univ-amu.fr/ihmt-moco.

## Methods

### Population and MRI acquisitions

Imaging experiments were performed on a 1.5T MRI system (Avanto, Siemens, Erlangen, Germany) with body coil transmission and a 32-channel receive-only head coil. Written informed consent was obtained from all subjects. Two groups of subjects were investigated: 6 healthy volunteers (5:1 women:men; mean age 28 years spanning from 21 to 44 years) and 7 patients with relapsing-remitting MS (6:1 women/men; mean age 32 years spanning from 21 to 51 years, mean disease duration = 72±84 months, median Expanded Disability Status Scale (EDSS) score=1.07, range=[0;3.5]). Subjects were instructed to remain still during the imaging protocol.

A whole-brain, slab-selective axial 3D-cartesian steady-state boosted-ihMT-GRE sequence was used with the optimized configuration described in (11): a pulsed MT preparation consisting in 12 Hanning-shaped pulses (0.5-ms long) repeated every 1 millisecond with frequency-alternation for dual-saturation, was interleaved with a segmented GRE readout (9 segments per readout train, readout (RO) flip angle of 7°, TR_RO_/TE=6.2/3.0 ms, and a receiver bandwidth of 370 Hz/voxel). The scheme of MT preparation-segmented GRE readout was repeated every TR=67.9 ms to acquire 3D MTw images (matrix size of 128×100×80) at a nominal 2 mm isotropic resolution. The effective RF saturation power over TR was B_1,RMS_=5.5 μT accounting for the partial Fourier saturation technique described in Ref. (11)). The images required to generate the parametric ihMTR maps were sequentially acquired and consisted of a reference image (MT_0_, no RF saturation pulse applied), followed by four MTw images obtained with RF saturation at positive frequency-offset (+8 kHz; MT_+_), dual frequency-offset (+/−8 kHz; MT_+/−_), negative frequency-offset (−8 kHz; MT_−_) and dual frequency-offset (−/+8 kHz MT_−/+_) respectively. No coil-based B_1_^−^ compensation from the constructor routines were applied on reconstructed data. To correct for Gibbs-ringing artefacts, the raw data were apodized using an isotropic 3D-cosine kernel.

### Multi-Contrast Motion Correction (MC-MoCo)

The proposed spatial motion correction method, called MC-MoCo thereafter, was compared to the freely available Motion Correction algorithm from the FMRIB’s Linear Image Registration Tool (MCFLIRT) (4) from the FSL package (version 5.0.8; available at: https://fsl.fmrib.ox.ac.uk/), using the averaging option for target building.

Both MC-MoCo and MCFLIRT rely on rigid-body transformations (6 degrees of liberty) since no non-linear deformations are expected between images based on the same readout in a single subject. Since native images (M_0_, MT_+_, MT_+/−_, MT_−_ and MT_−/+_) have distinct contrasts, a mutual information metric (12) was systematically employed, and images were linearly resampled upon transformation.

Figure 1 illustrates the proposed MC-MoCo method. The whole process relies on functionalities provided in the Advanced Normalization Tools (ANTs) registration package (5) (version 2.1.0; available at: http://stnava.github.io/ANTs/) and can be summarized as follows: (i) Nick’s nonparametric nonuniform normalization (N4) bias-field correction (13) and skull-stripping of the native images, (ii) target building based of the MTw images, (iii) registration of each native skull-stripped images onto the targset, (iv) application of the estimated transformations to each native image.

**Figure 1.**
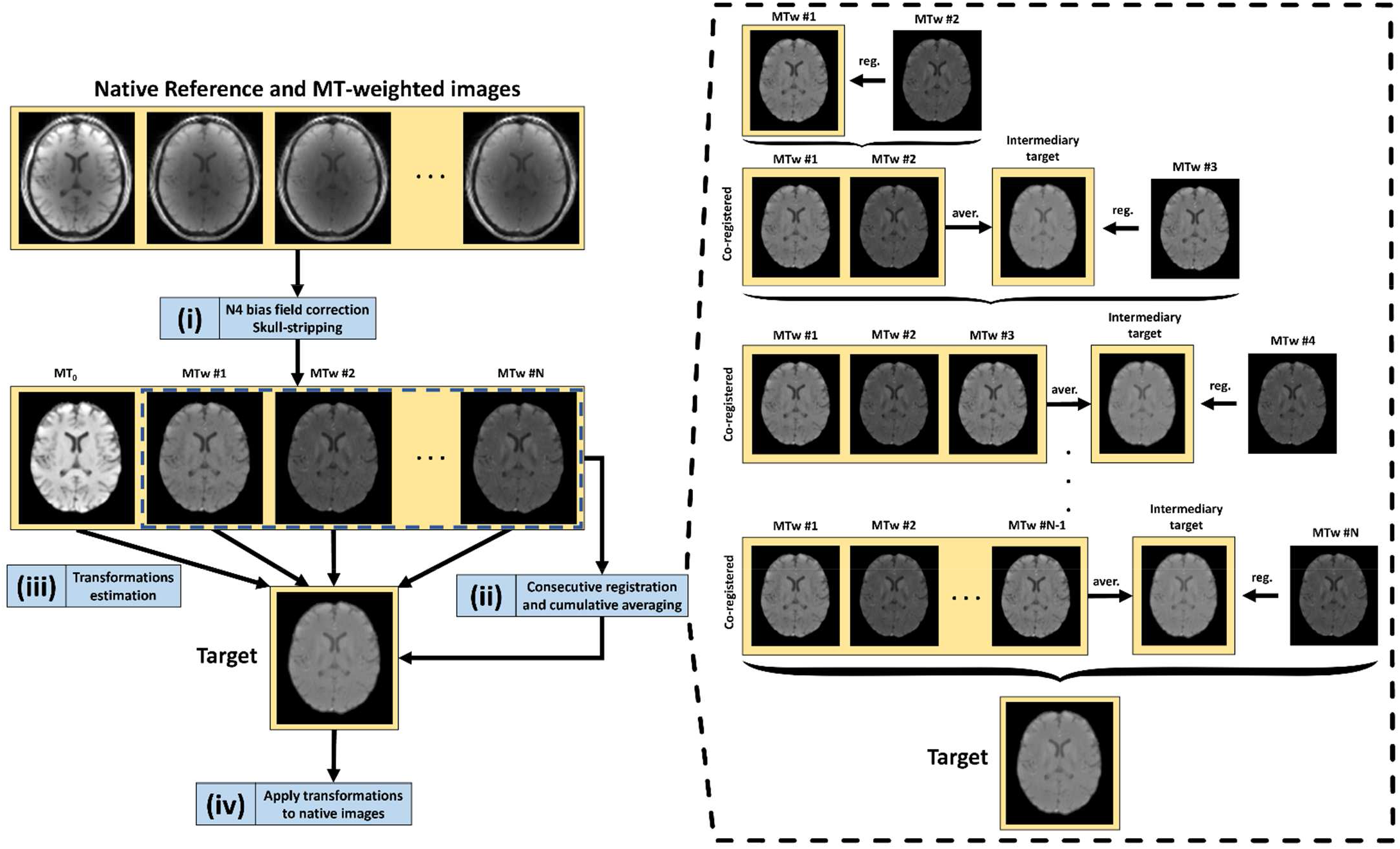
Schematic depiction of the proposed motion-correction algorithm. A first processing step consisting in a bias field correction and a skull-stripping is performed over all individual native images (i). MT-weighted images are thereafter used to build a robust target in a progressive cumulative-averaging framework, in which a consistency between co-registered images is perpetuated. Finally, transformations of all individual images onto the newly built target are estimated (iii), and applied to native images (iv). All transformations are rigid and estimated using the mutual information metric.

A rationale for steps (i) and (ii) is given:

i. In order to enhance the sensitivity of the transformation estimation to brain structures, a skull-stripping step was first performed. This allowed minimizing contributions of out-of-brain features (e.g. various signal intensities in the scalp across MTw and reference images) throughout the registration process. The skull-stripping procedure was realized with the *antsBrainExtraction.sh* routine (14) applied on in-house templates accounting for the three subsequent contrasts (namely from reference, single- and dual-saturation images). The bias field correction step helped enhance image contrasts for skull-stripping purpose.
ii. In order to build a high-SNR and robust target for co-registration, MTw images were co-registered in a cumulative-averaging manner, as depicted in the right-side of Figure 1. Co-registration of similarly contrasted images further limits registration errors, while allowing for an independent and trustworthy target building.

### Motion evaluation

In order to assess the efficiency of both MCFLIRT and MC-MoCo methods to correct for motion, four sets of landmarks were positioned at anatomical regions over each image. These landmarks were defined at spatial positions demonstrating a fair sensitivity to movements: (i) fissure in the right superior frontal gyrus (R-SFG), (ii) fissure in the right precentral gyrus (R-PCG), (iii) fissure in the left middle frontal gyrus (L-MFG) and (iv) upper (bottom-top direction) space between right and left subcallosal cingulate gyrus (LR-SCCG). Landmarks were placed by an expert in neurology (J.P.) on the native images and MCFLIRT and MC-MoCo images using the Markups plugin from 3D Slicer (version 4.10.2; available at: https://www.slicer.org/) (15). Representative positionings of these landmarks are depicted in Figure 2 on 3D rendered brain images (16).

**Figure 2:**
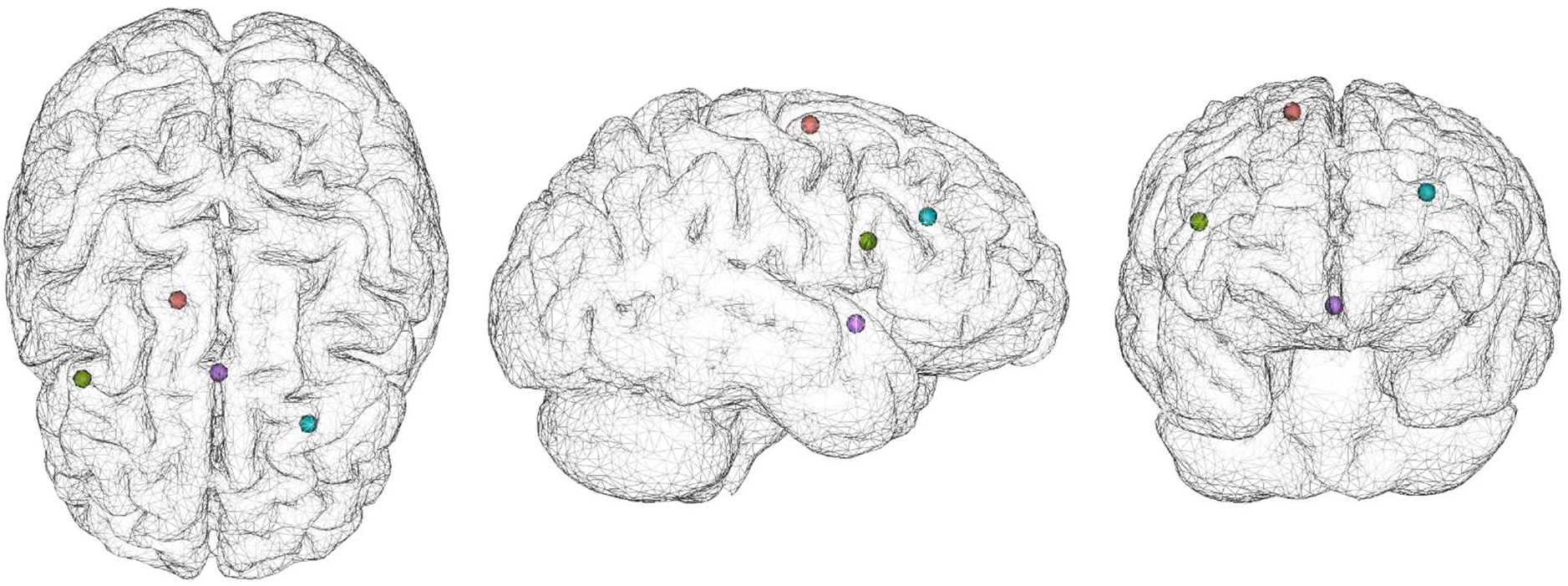
Overlaid representative spatial positionings of the selected landmarks on a 3D-rendered brain (red: R-SFG; green: R-PCG; blue: L-MFG; purple: LR-SCCG) over top-bottom view (left), right-left view (middle) and frontal-occipital view (right).

**Figure 3:**
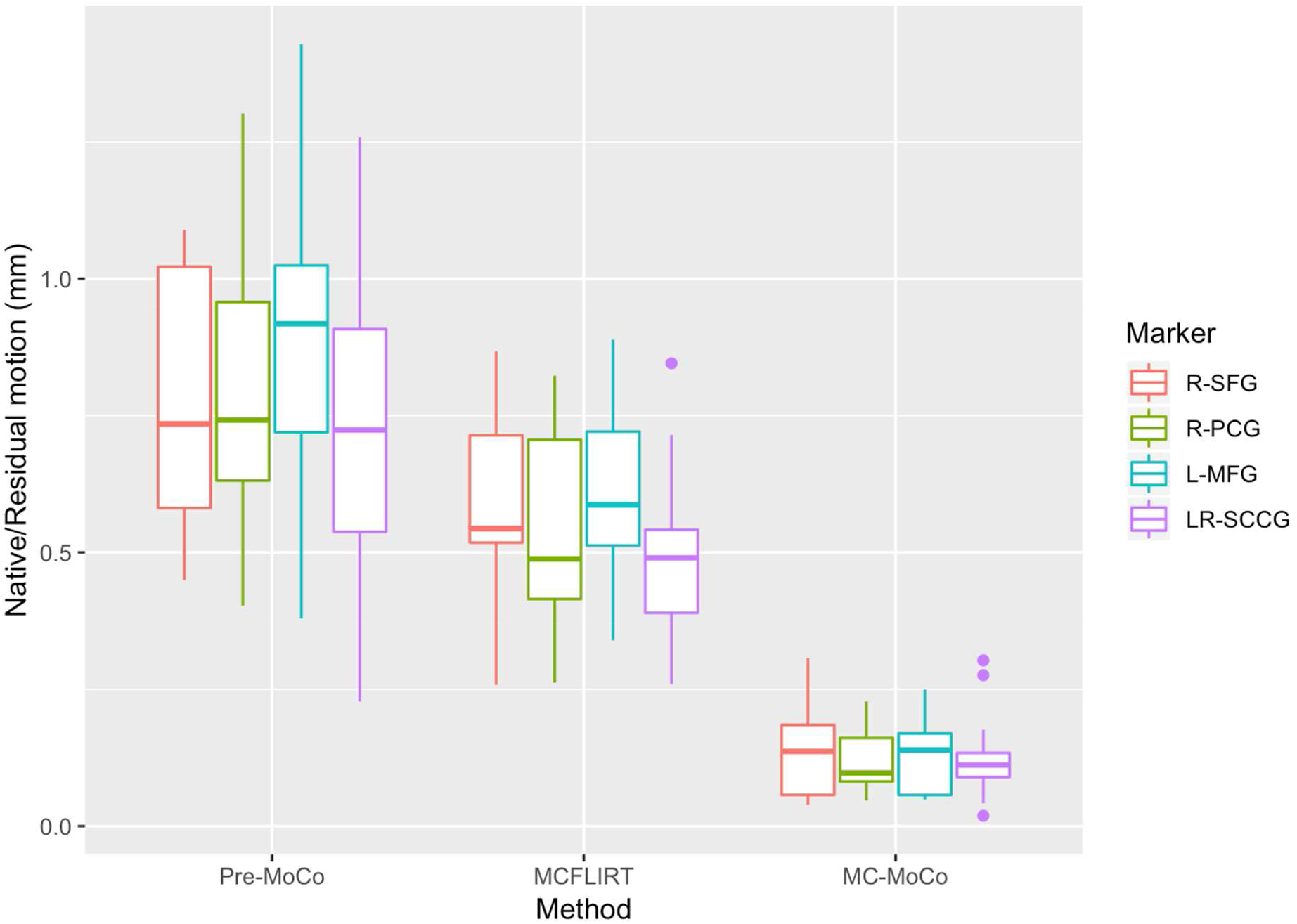
Boxplots of native motion (left), measured on native non-corrected images and residual motion measured on MCFLIRT (middle) and MC-MoCo (right) corrected images across the four markers for all subjects.

For each subject and each set of images (uncorrected, MCFLIRT- and MC-MoCo-corrected), a reference point of each landmark was defined as the centroid of the landmark locations across the images of the set. The reference point was used to quantify the amount of native motion (uncorrected images) and residual motion (MCFLIRT- and MC-MoCo-corrected images) by averaging the distances between the reference point and its corresponding landmarks over all the subjects. The residual motion reflects the quality of the motion-correction algorithm: a better algorithm will exhibit less residual motion, i.e. a lower distance between a landmark and its reference point. Additionally, for an unbiased algorithm, this criterion should be consistent for every landmark and for both groups of subjects (healthy volunteers and MS patients).

### Statistical analyses

Based on the previously-defined motion quantification, statistical analyses were performed to evaluate the performance of MC-MoCo and MCFLIRT, and to verify that the behavior of both methods was constant across landmarks and between the two groups of subjects. For these analyses, we used a mixed-effect linear model with fixed effects for the motion correction method, the landmark, and the subject group, and with a random effect for the subject to account for the non-independence of the observations within a subject. Shapiro-Wilk and Mann-Whitney tests were used to test for normality and median difference of distributions, respectively. The analysis code based on R is available at https://github.com/lamyj/mcmoco-data (commit 9e0555b was used for the presented results).

## Results

A boxplot of the averaged (over the subjects) native motion evaluated on the images before motion correction and residual motion evaluated after application of MCFLIRT and MC-MoCo is shown in Figure 3 and indicates a lower residual motion for MC-MoCo compared to that of MCFLIRT for all landmarks. The residual motion value was additionally rather constant across all markers for MC-MoCo.

The residuals of the mixed-effects linear model showed neither deviation from a normal distribution (Shapiro-Wilk test, p=0.57) nor apparent structure in the variance of the residuals, thus validating the model. The analysis of the individual factors shows a significant effect of the motion-correction methods (p<0.01), indicating a difference between the non-corrected images and the motion-corrected images, but no difference either across the landmarks (p=0.14) or between the two groups of subjects (p=0.12), thereby indicating that differences in residual motions depends only on the algorithm. Further linear hypotheses tests comparing the native motion from residual motions derived from MCFLIRT and MC-MoCo show reductions of −0.25 mm (p<0.01) for MCFLIRT and −0.68 mm (p<0.01) for MC-MoCo. Additionally, reductions of the residual motion of MC-MoCo and MCFLIRT were statistically different (p<0.01). P-values were corrected for multiple comparisons.

Figure 4 illustrates native and corrected motion over a sagittal magnified region for a subject presenting a motion mainly characterized by a head lift-up. Voxel re-alignment can be observed following both motion correction methods, with a qualitatively better result with the MC-MoCo technique compared to MCFLIRT. Representative axial, sagittal and coronal views of MT_0_ and MTw images for all subjects before and after corrections are provided as animated images in Supporting Information. To enhance visibility, images were bias-field corrected, Laplacian-sharpened, and intensities were normalized.

**Figure 4:**
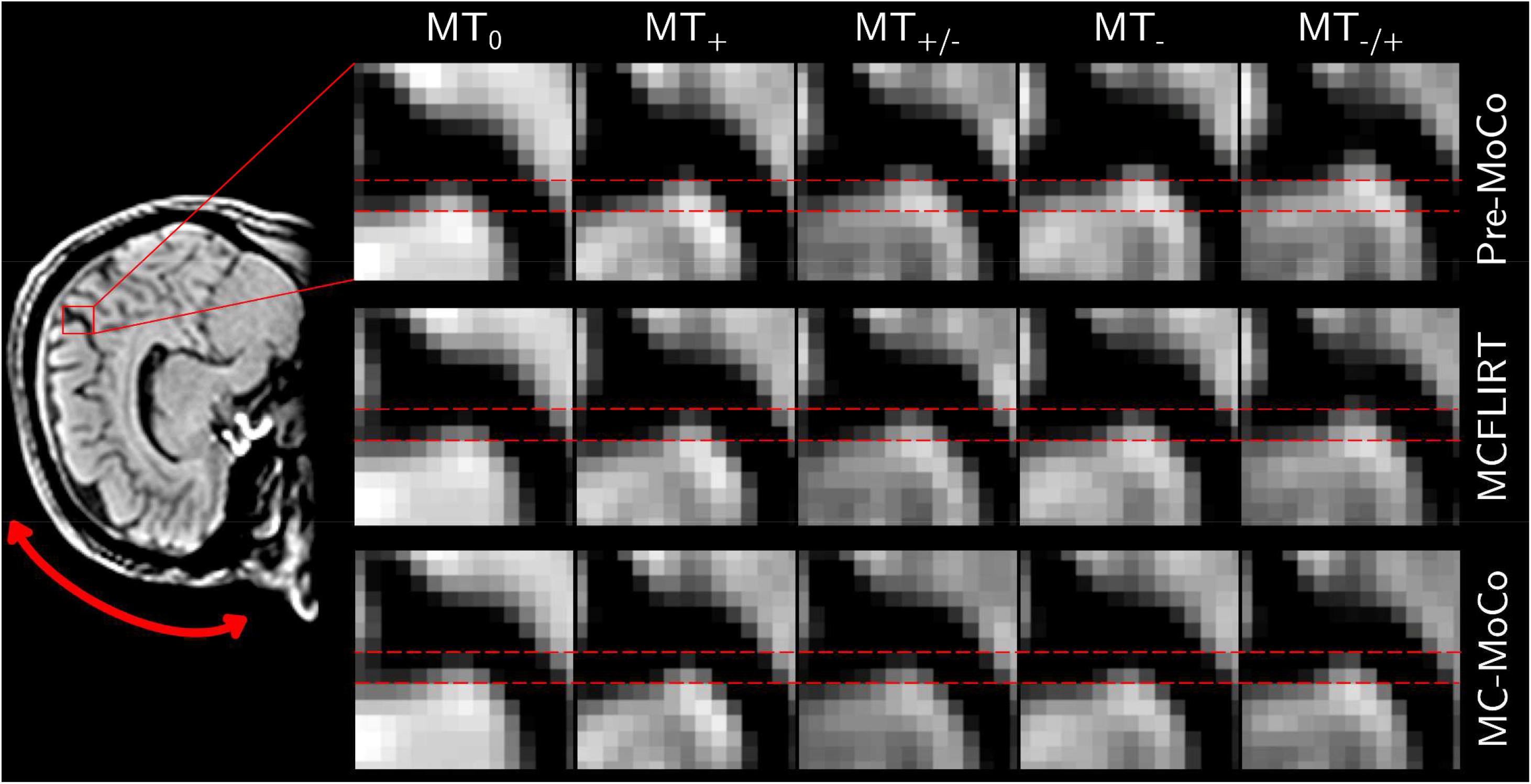
Representative magnified region over reference and weighted images before (left column) and after application of MCFLIRT (middle column) and MC-MoCo (right column). The qualitative main direction of motion of the selected subject consisted of a head lift-up, depicted with the double-headed red arrow. Dashed red lines are provided as focusing features to better apprehend native and residual motions.

Figure 5a shows exemplary orthogonal views of ihMTR maps before and after motion correction. Following MCFLIRT and MC-MoCo applications, severe artefacts related to misalignment upon image combination (red arrows for brain-peripheral artefacts and blue arrows for peri-ventricular ones) are removed, with a better performance using the proposed MC-MoCo method. Finally, Figure 5b depicts the effect of motion correction in an exemplary MS lesion where a qualitative superior sharpness is reached following the application of MC-MoCo. Figure 5c shows the lesion distributions of the ihMTR image difference between MC-MoCo and Pre-MoCo, and MCFLIRT and Pre-MoCo, emphasizing that MCFLIRT yields a minor improvement (narrow distribution) compared to MC-MoCo (wide and bimodal distribution). A significant median difference of the voxel distributions in the lesion is observed between Pre-MoCo and MC-MoCo (Mann-Whitney U-test; p=0.001), but not between Pre-MoCo and MCFLIRT (p=0.73).

**Figure 5:**
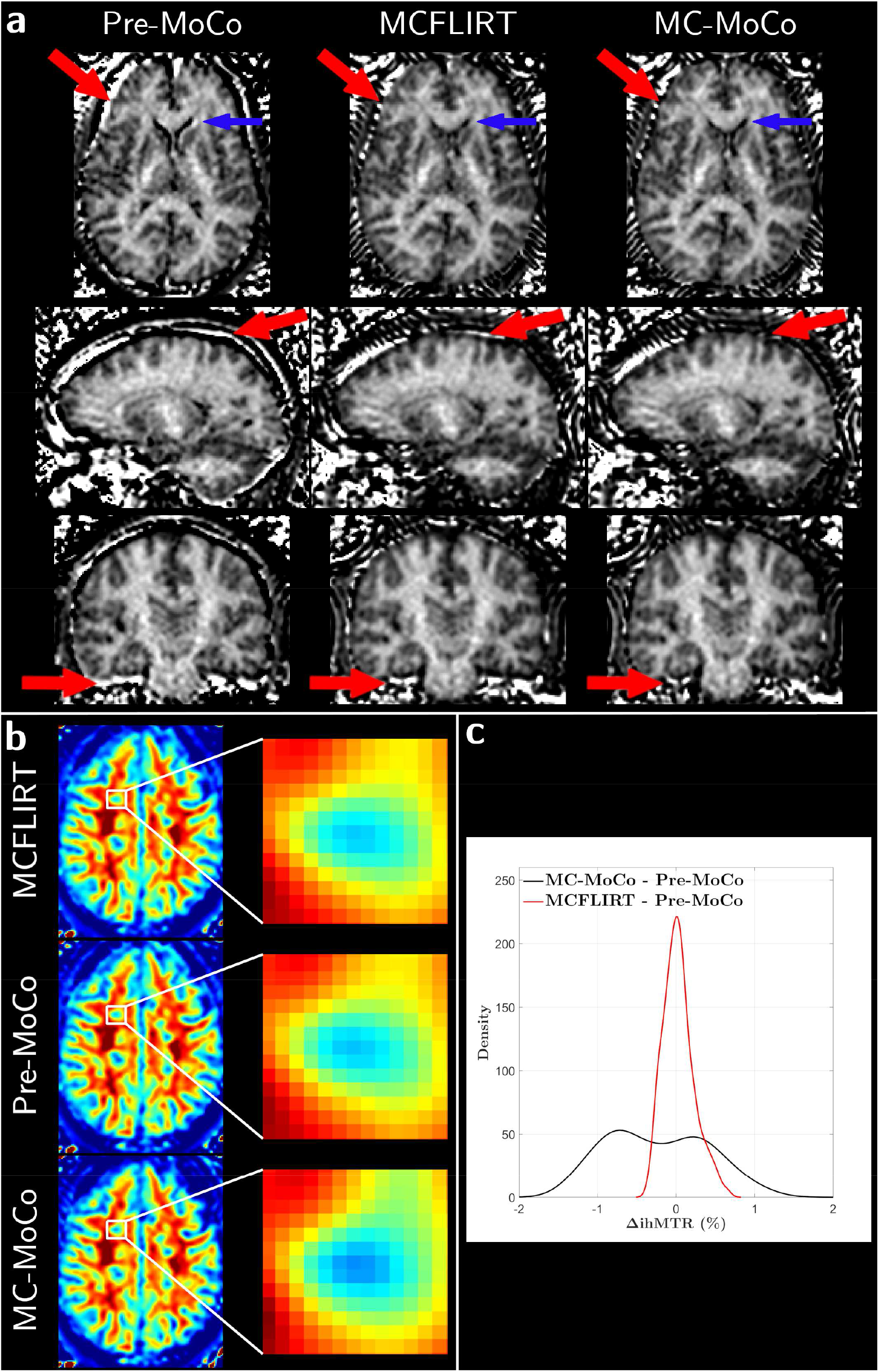
**a**: Representative orthogonal views of ihMTR maps from a single subject generated before (left column) and after motion correction using MCFLIRT (middle column) and MC-MoCo (right column). Red and blue arrows indicate remarkable regions showing misalignment-related artefacts detected in the Pre-MoCo ihMTR map. **b**: Representative ihMTR MS lesion before and after motion correction. **c**: Distributions of the image difference (ΔihMTR) between MC-MoCo and Pre-MoCo, and MCFLIRT and Pre-MoCo about the selected lesion.

## Discussion

We developed a refined motion-correction algorithm adapted for MRI techniques whose metrics of interest rely on the combination of a small number of images with various contrasts, and evaluated it in the particular case of the ihMT MRI technique.

The key point of the MC-MoCo method lies in the target building step. As opposed to other motion-correction methods based either on an averaged target over all native images or on a single selected reference image, the target is built using an iterative and updated pattern, which provides a reliable result given a limited number of images. A readily usable alternative for an iterative template building is provided using tuned options by ANTs (in *antsMultivariateTemplateConstruction2.sh* as of ANTs version 2.2.0). Nevertheless, using the proposed method, the residual motion after correction was significantly decreased compared to MCFLIRT.

The comparison of MC-MoCo and MCFLIRT was performed at a global level to emphasize that both methods are usable out-of-the-box. This comparison may induce a bias, as no image enhancement (e.g. skull-stripping or bias-field correction) is natively included in MCFLIRT. Moreover, the MC-MoCo pipeline can be implemented based on individual features from packages other than ANTs (e.g. SPM, FSL and AFNI (17)), although slight differences in results should be expected since the implementation details of each package are intrinsically different. Although each method and all relevant steps may be further optimized, the presented results provide a fair comparison from the perspective of the end-user.

In our investigation, the MCFLIRT algorithm was tuned to build a target by averaging all available images acquired for a single subject as it is done for standard functional MRI motion correction procedures to yield a high-SNR target. This approach had the advantage of avoiding to arbitrarily select a reliable reference image among all available contrasts, themselves possibly varying with sequence parametrization. Averaging hence results to an objective single choice for target building.

Only rigid registrations were considered throughout the correction processes of the MC-MoCo method. These transformations contributed to maintain original geometries, which is an important asset for investigating pathological tissue featured by modification of structure (e.g. MS lesions). Note that in the case of non-conventional acquisition schemes (e.g. echo planar imaging), potential non-linear image distortions limit the use of the rigid transformation.

The skull-stripping step in MC-MoCo is part of the image enhancement process prior to the registration steps and was performed using a template-based framework. Alternatively, the Brain Extraction Tool (BET) (18) in FSL may be considered as it is faster than a template-based approach and readily usable, but unfortunately necessitates subject-wise parameter tuning in order to achieve correct results on every image.

No significant group effect (control versus MS patients) was found on the estimated model. The MC-MoCo method is therefore equivalently efficient for both populations. Given the proposed framework for re-alignment, the MC-MoCo method is expected to yield satisfying results in the case of agitated subjects, as long as images are not affected by intra-scan movement artefacts. Nonetheless, further investigations have to be performed to assess its robustness in the case of severe motion.

The proposed MC-MoCo method could be readily applied and tested to other imaging modalities whose final contrast generation share similar features with ihMT (e.g. ASL, CEST or MT imaging). Indeed, all these techniques include i) a reconstruction based on a combination of a small number of raw images, ii) a range of varying image contrast and SNR across experiments and iii) a sensitivity to misalignment upon images combination.

In this study, images were acquired without coil-based B_1_^−^ bias correction in order to prevent spatial intensity inconsistencies and local noise amplification. Hence, motion correction will most likely re-align voxels presenting different B_1_^−^ sensitivity, resulting in discrepancies in terms of actual voxel-wise signal intensity. We acknowledge that this may foster bias in computationally derived maps. Nonetheless, this bias field is slowly varying spatially, and should not represent a critical issue if the subject motion remains reasonable. In addition, processing an increasing number of images may benefit in smoothing this effect.

Retrospective and prospective navigator-based methods have proven to be of great interest for motion correction (1, 19). However, prospective motion-correction may be incompatible with parametric imaging. For instance, in the ihMT-GRE technique, the magnetization is purposefully maintained in a steady-state, and RF-based navigator methods may disrupt magnetization, biasing further analyses. Thus, retrospective motion-correction methods such as the proposed MC-MoCo appear to be more suited for such applications.

## Conclusion

MC-MoCo is an efficient method for motion correction of the human brain in MRI techniques whose metrics of interest rely on the combination of a small number of images with various contrasts, such as ihMT. Image re-alignment further benefits registration processes between composite maps and anatomical images by restoring refined spatial features (e.g. MS lesions and cortical ribbon). Hence, MC-MoCo is a promising technique to explore the radiological aspects in MS, and can be routinely employed for the generation of ihMT-related maps.

## Supporting information

Supplemental Materiel

## Acknowledgements

This work was supported by the SATT Sud-Est (France), the French Association pour la Recherche sur la Sclérose En Plaques (ARSEP), Roche Research Foundation (Switzerland) and French National Research Agency, ANR [ANR-17-CE18-0030].

The authors thank V. Gimenez, P. Viout, L. Pini and C. Costes for technical support and management, as well as Dr. Fanny Munsch for the careful manuscript review.

## Supporting Information

Additional Supporting Information may be found in the online version of this article, presenting animated views of native and motion corrected images (both MC-MoCo and MCFLIRT) for each subject.

## Notes

### Competing Interest Statement

The authors have declared no competing interest.

