## Supplemental Materiel for "A Motion Correction Strategy for Multi-Contrast based 3D parametric imaging: Application to Inhomogeneous Magnetization Transfer (ihMT)"

### Slide 1
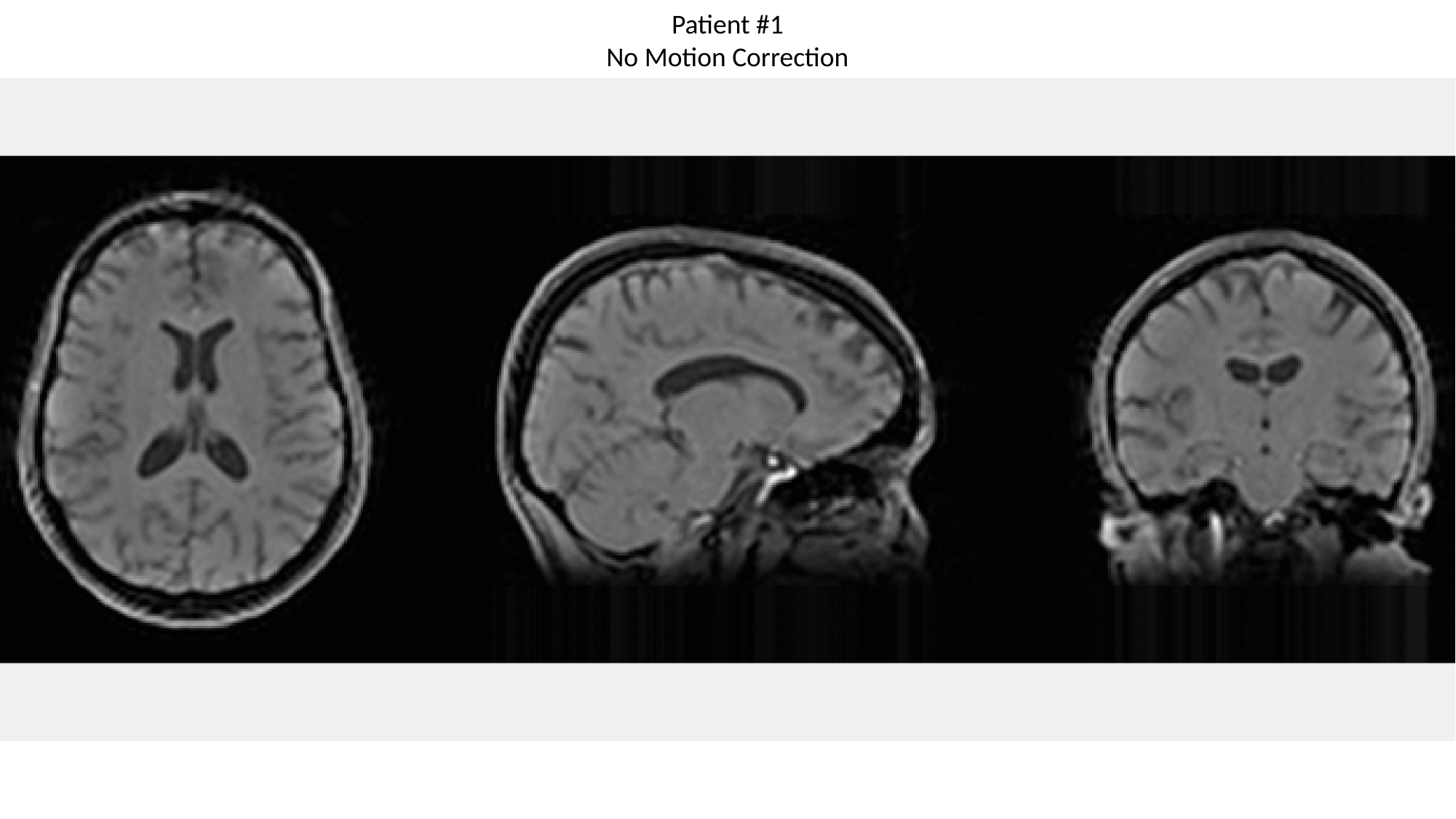

Patient #1
No Motion Correction

### Slide 2
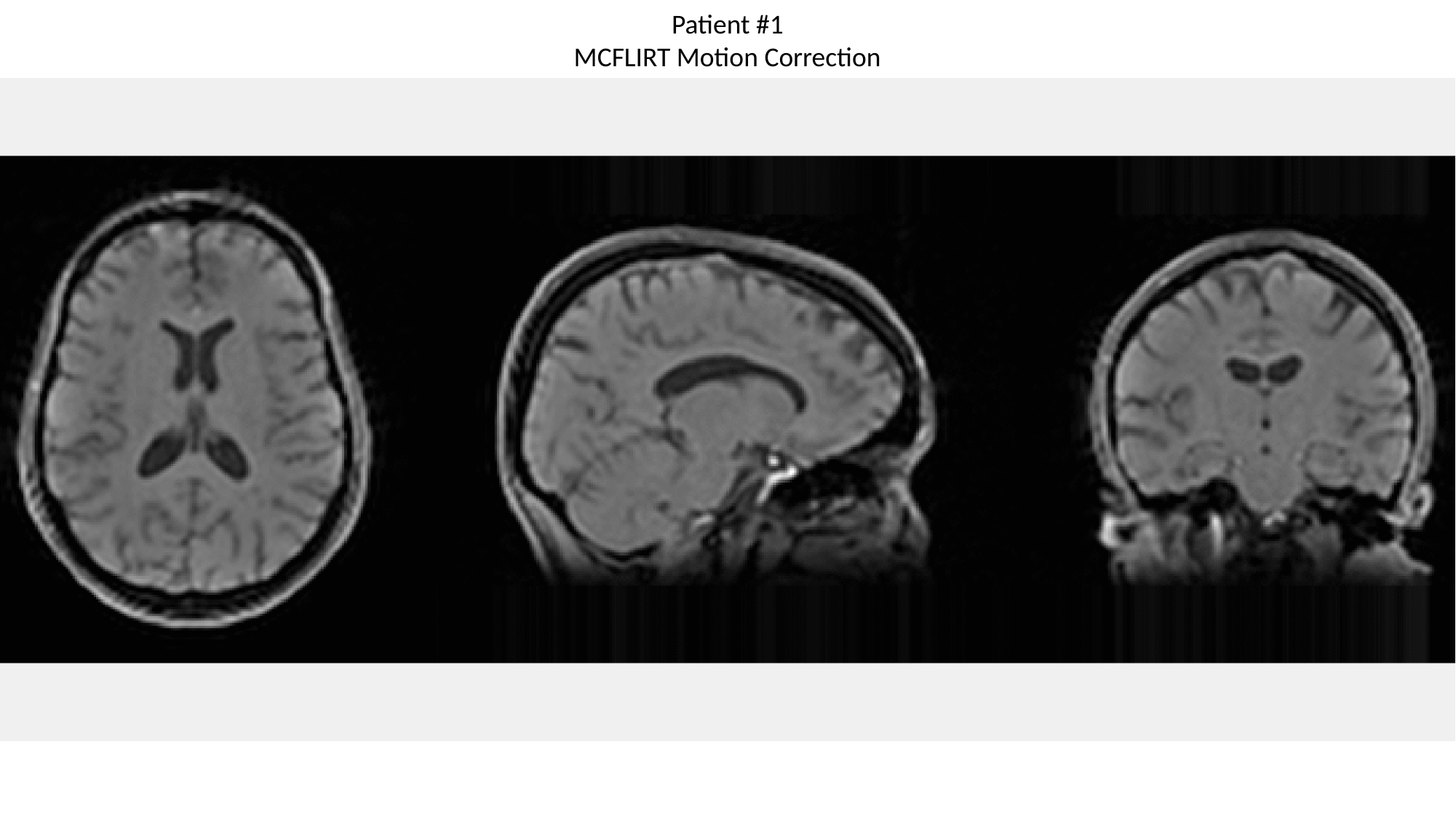

Patient #1
MCFLIRT Motion Correction

### Slide 3
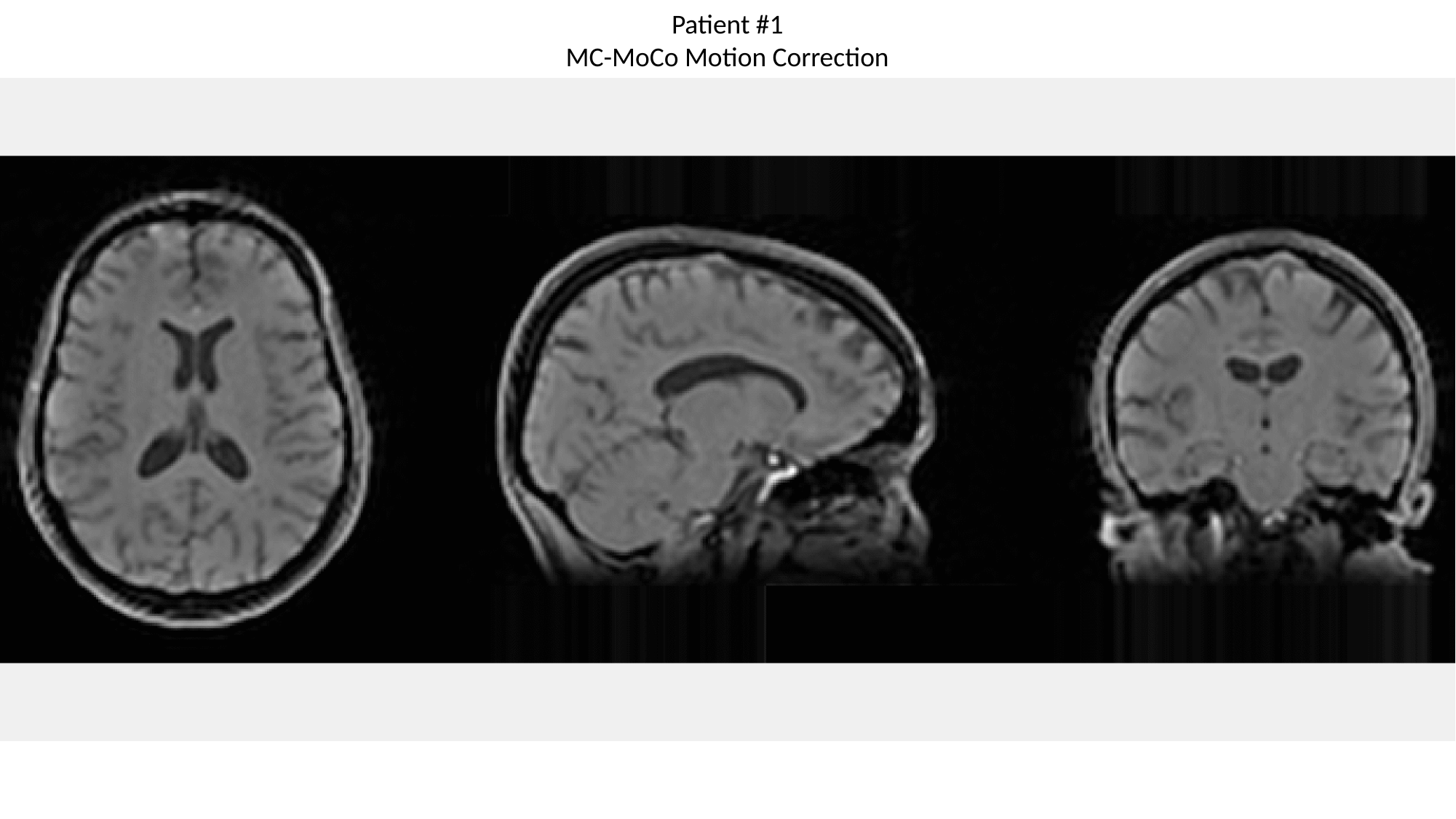

Patient #1
MC-MoCo Motion Correction

### Slide 4
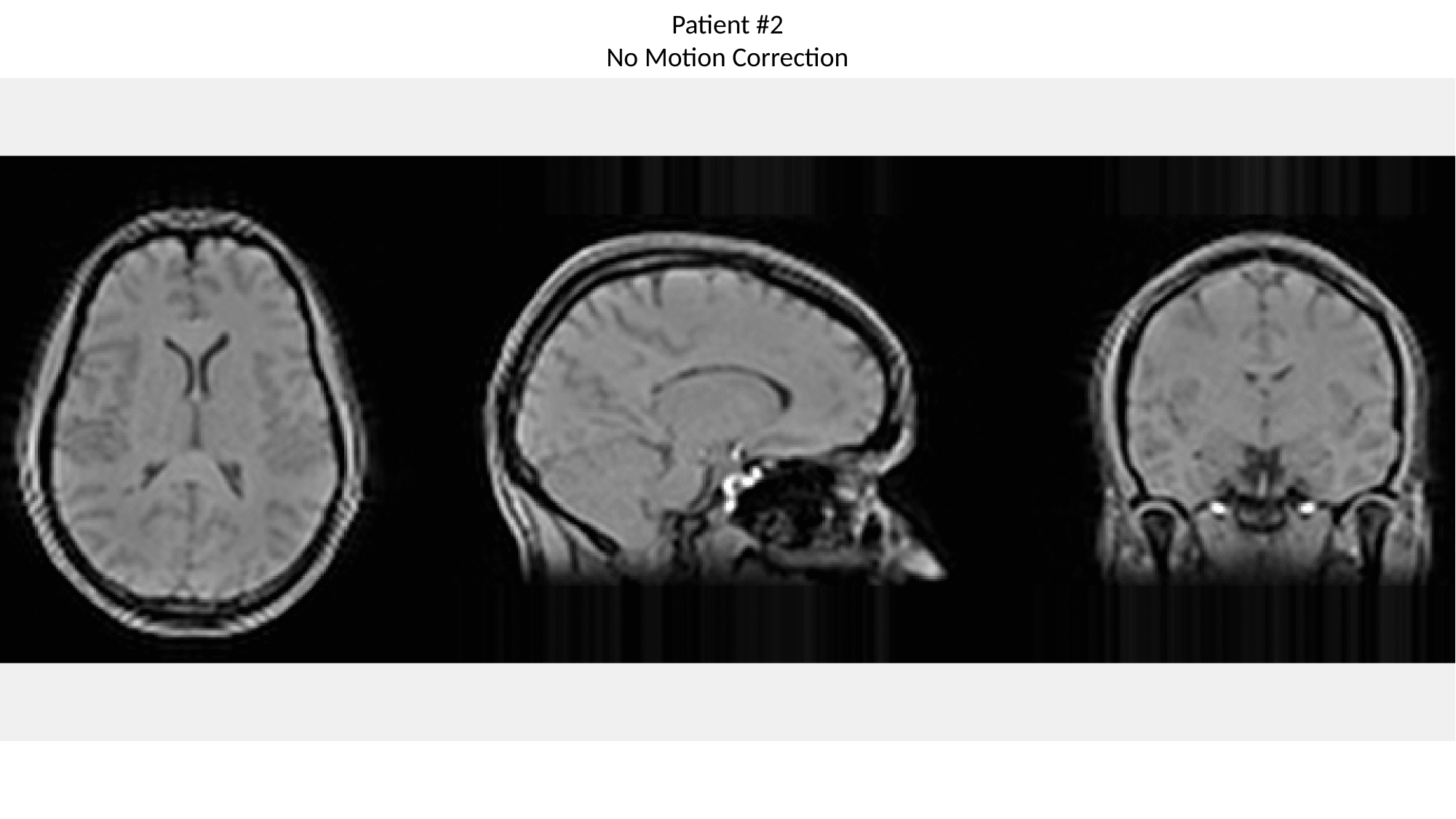

Patient #2
No Motion Correction

### Slide 5
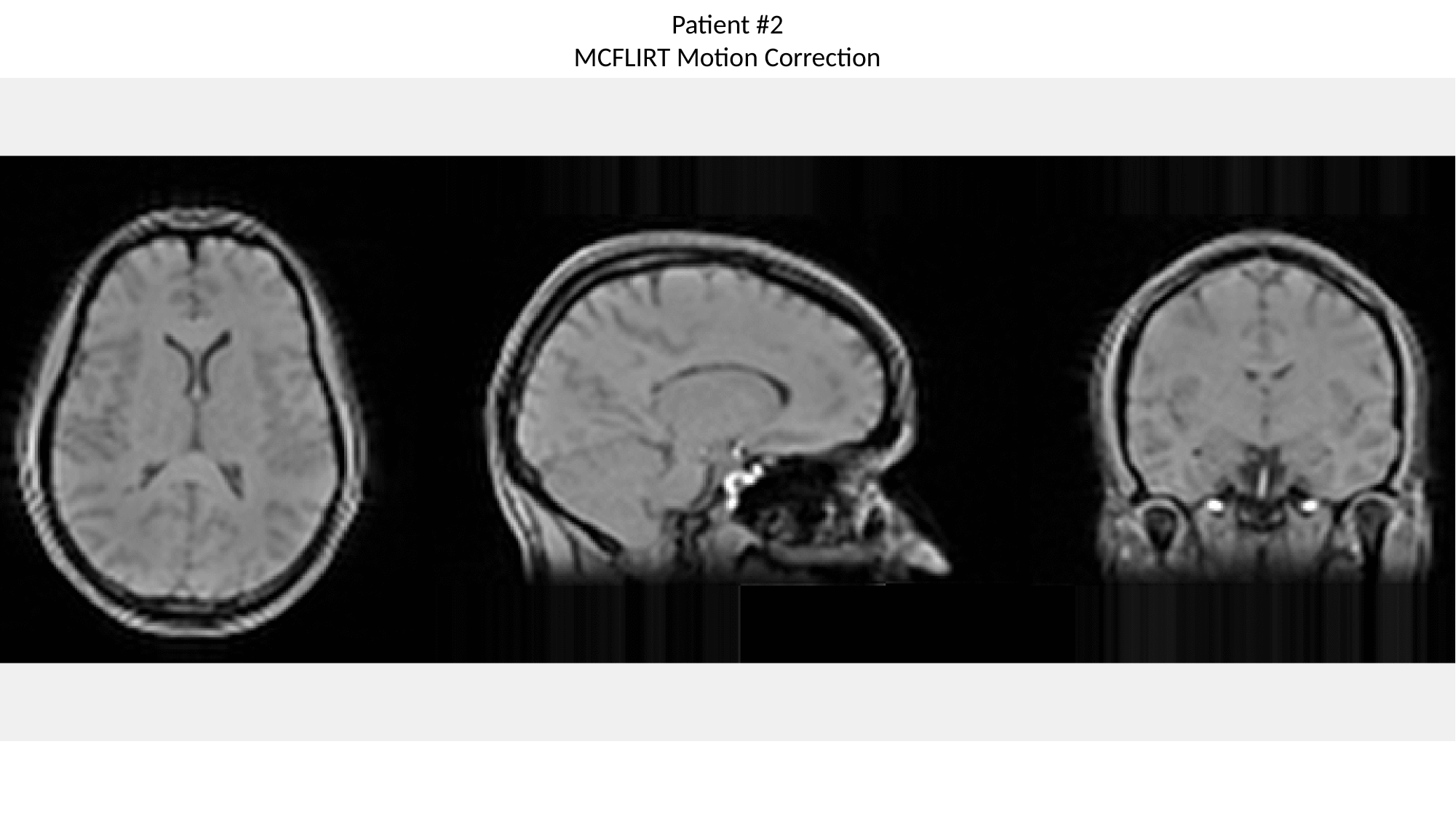

Patient #2
MCFLIRT Motion Correction

### Slide 6
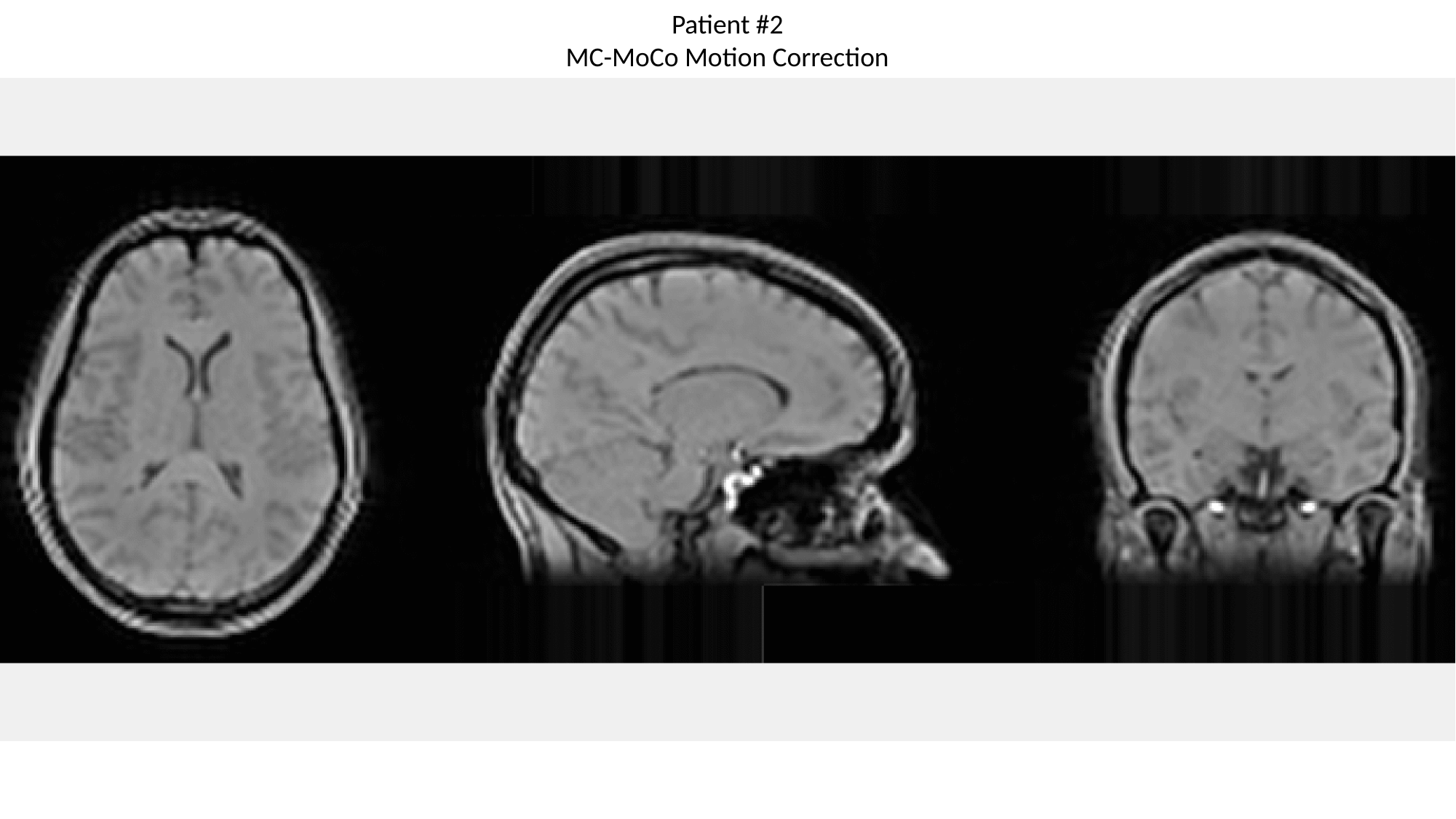

Patient #2
MC-MoCo Motion Correction

### Slide 7
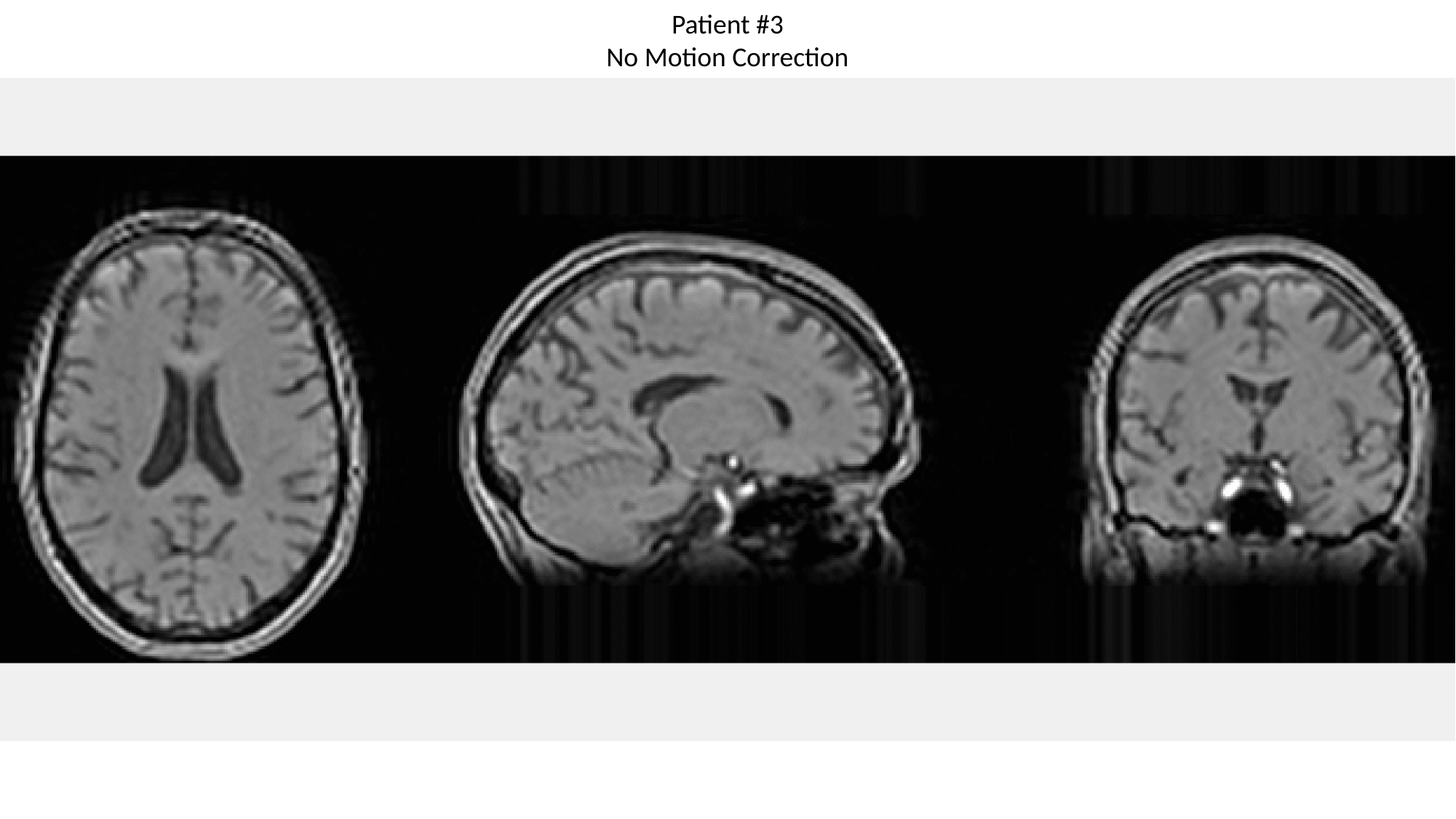

Patient #3
No Motion Correction

### Slide 8
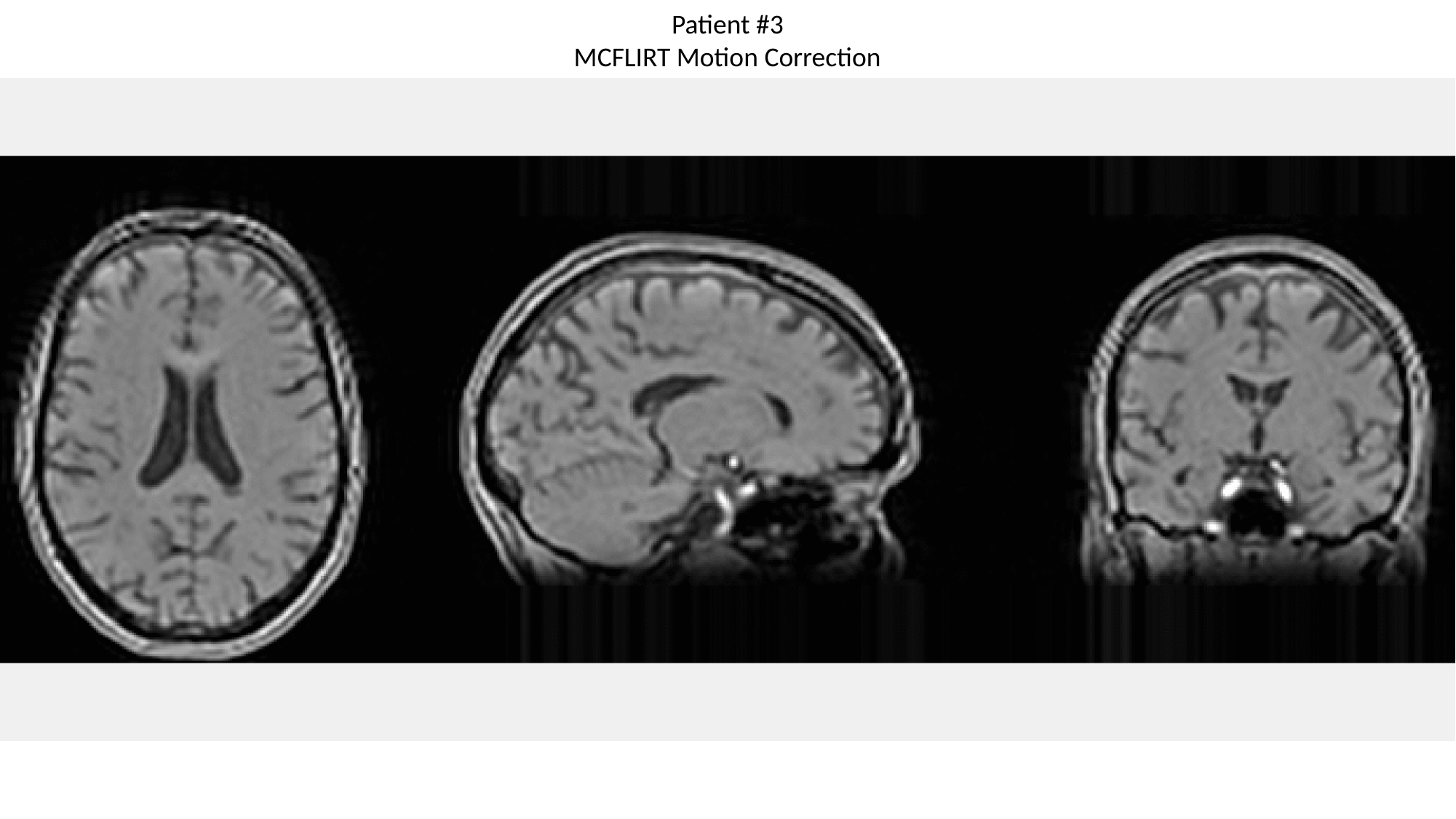

Patient #3
MCFLIRT Motion Correction

### Slide 9
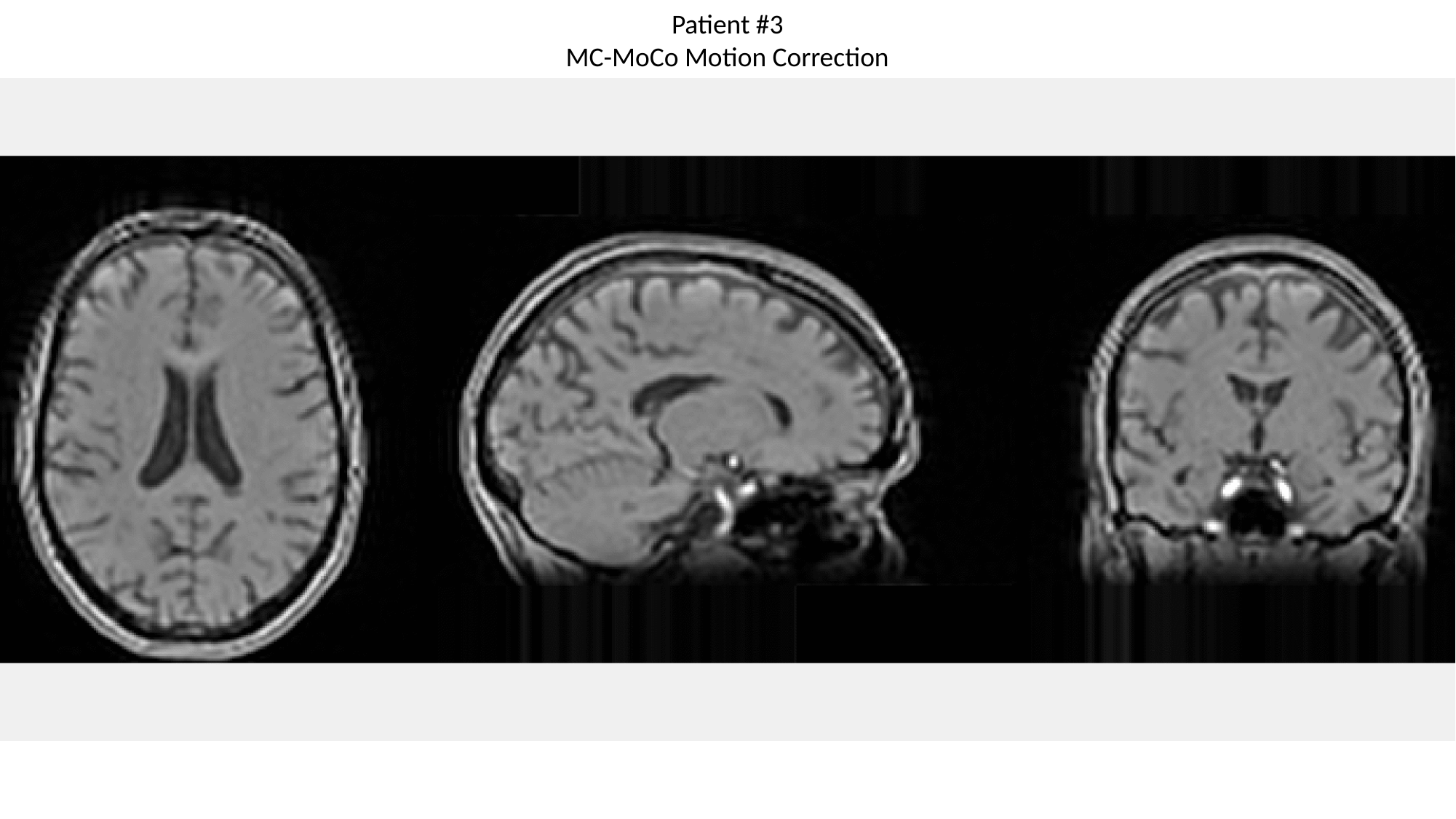

Patient #3
MC-MoCo Motion Correction

### Slide 10
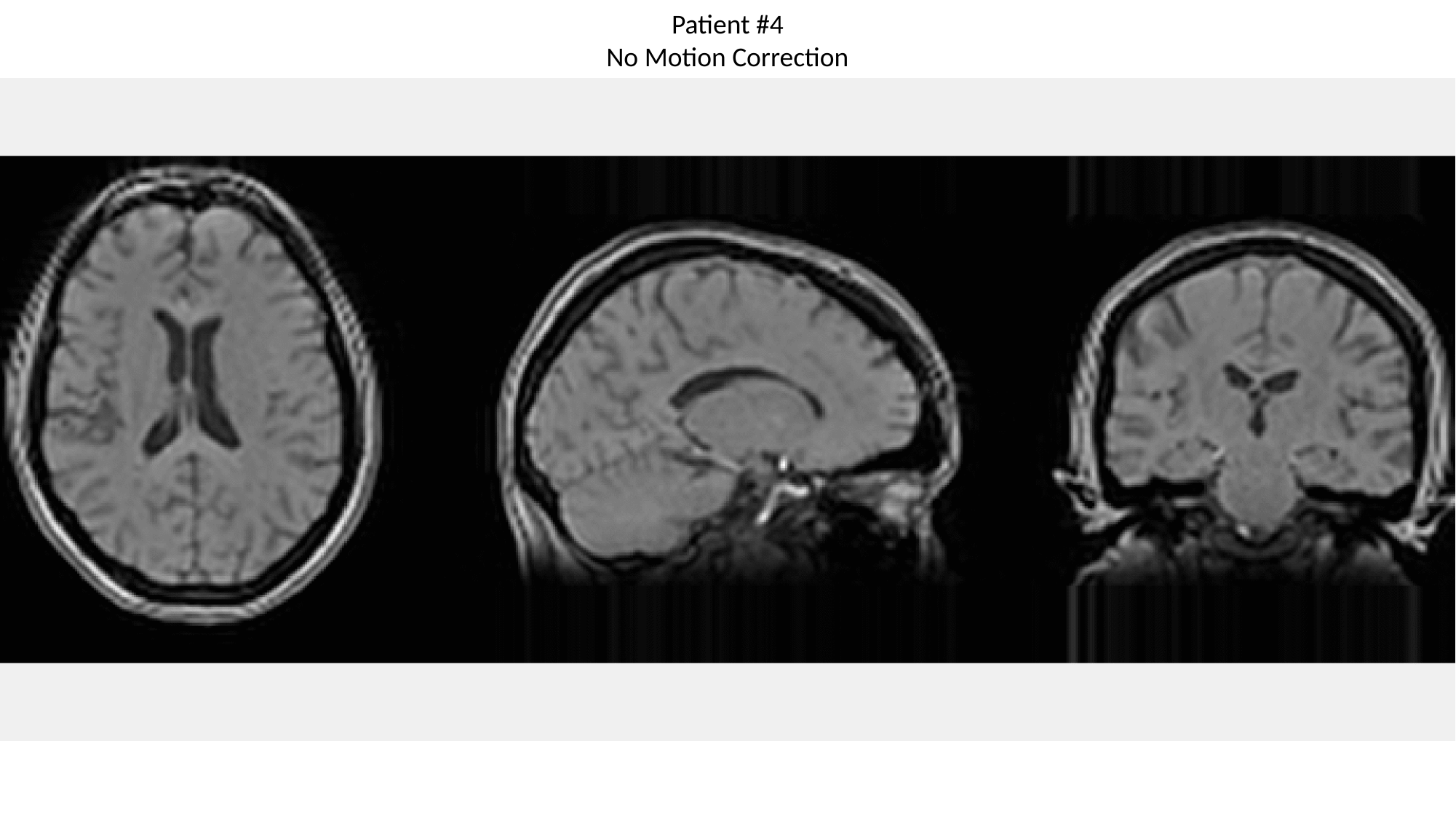

Patient #4
No Motion Correction

### Slide 11
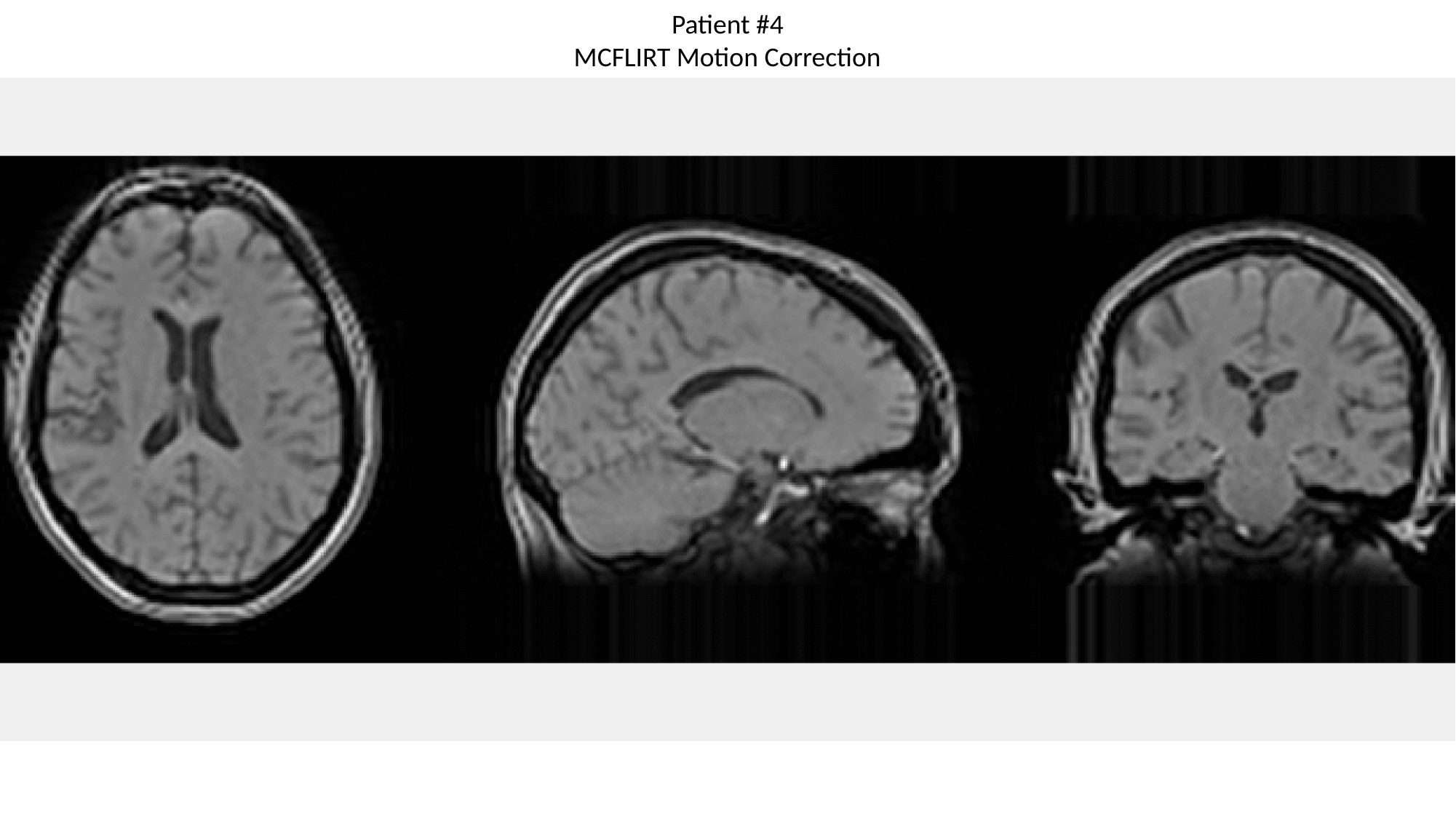

Patient #4
MCFLIRT Motion Correction

### Slide 12
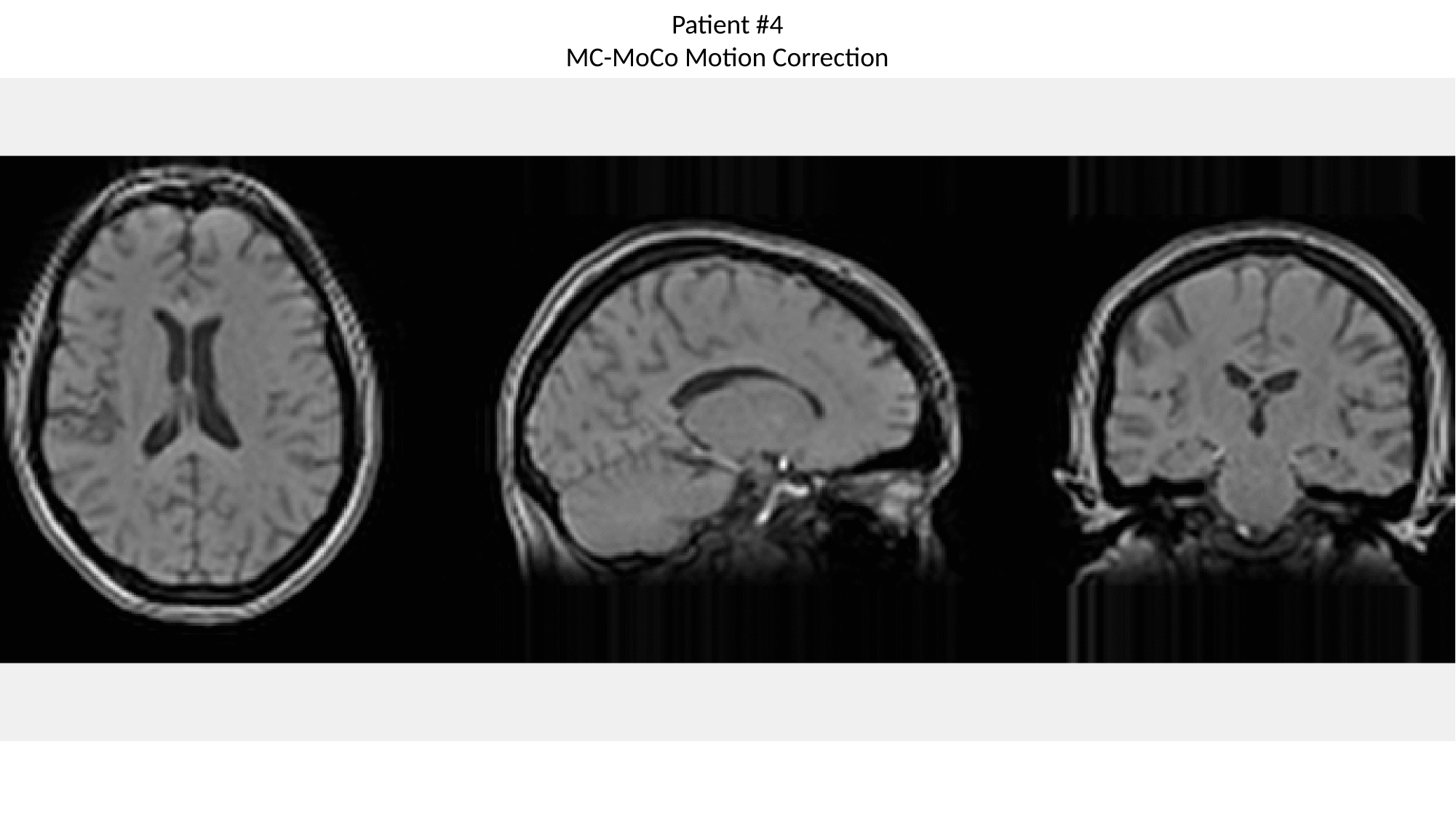

Patient #4
MC-MoCo Motion Correction

### Slide 13
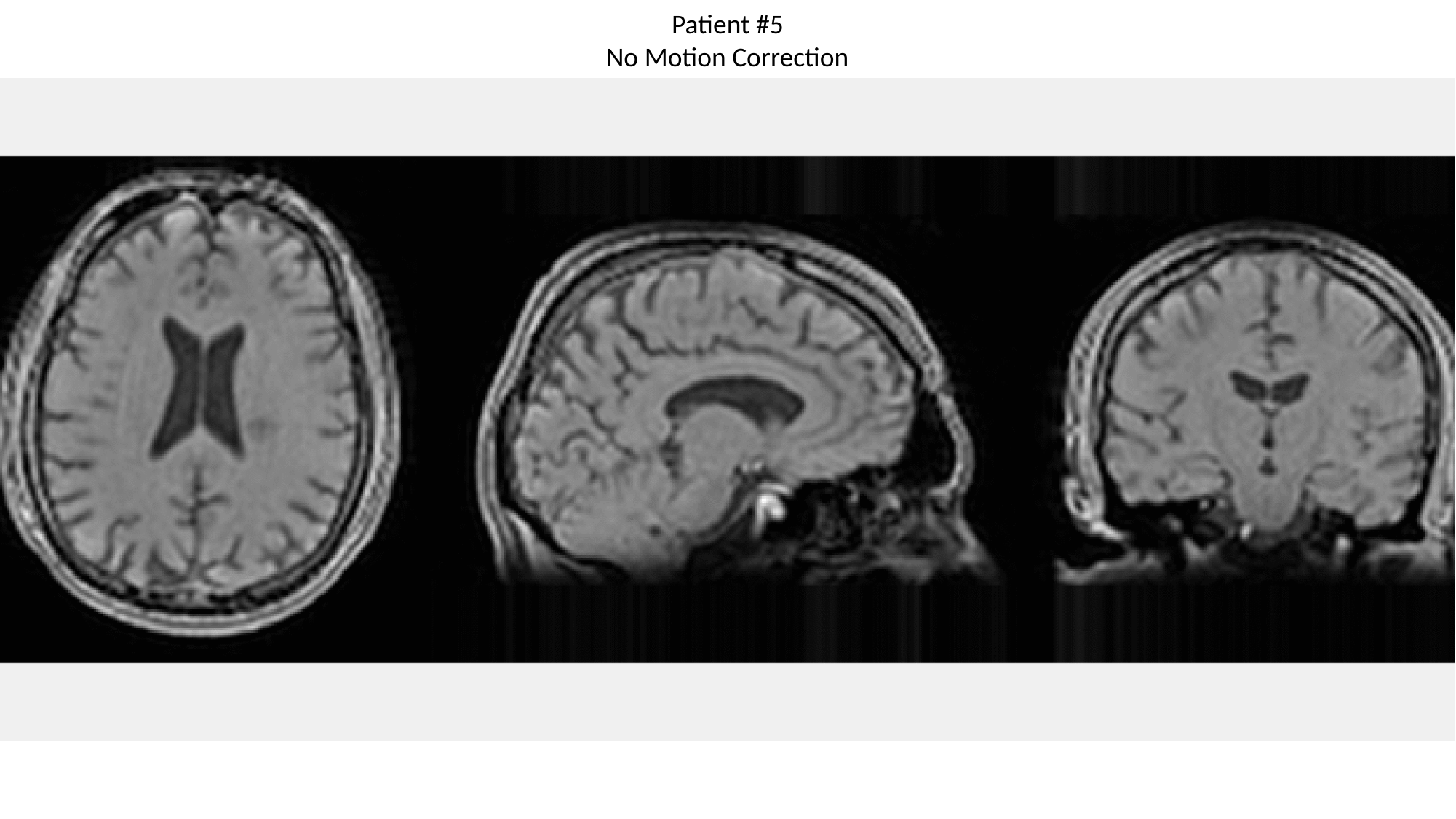

Patient #5
No Motion Correction

### Slide 14
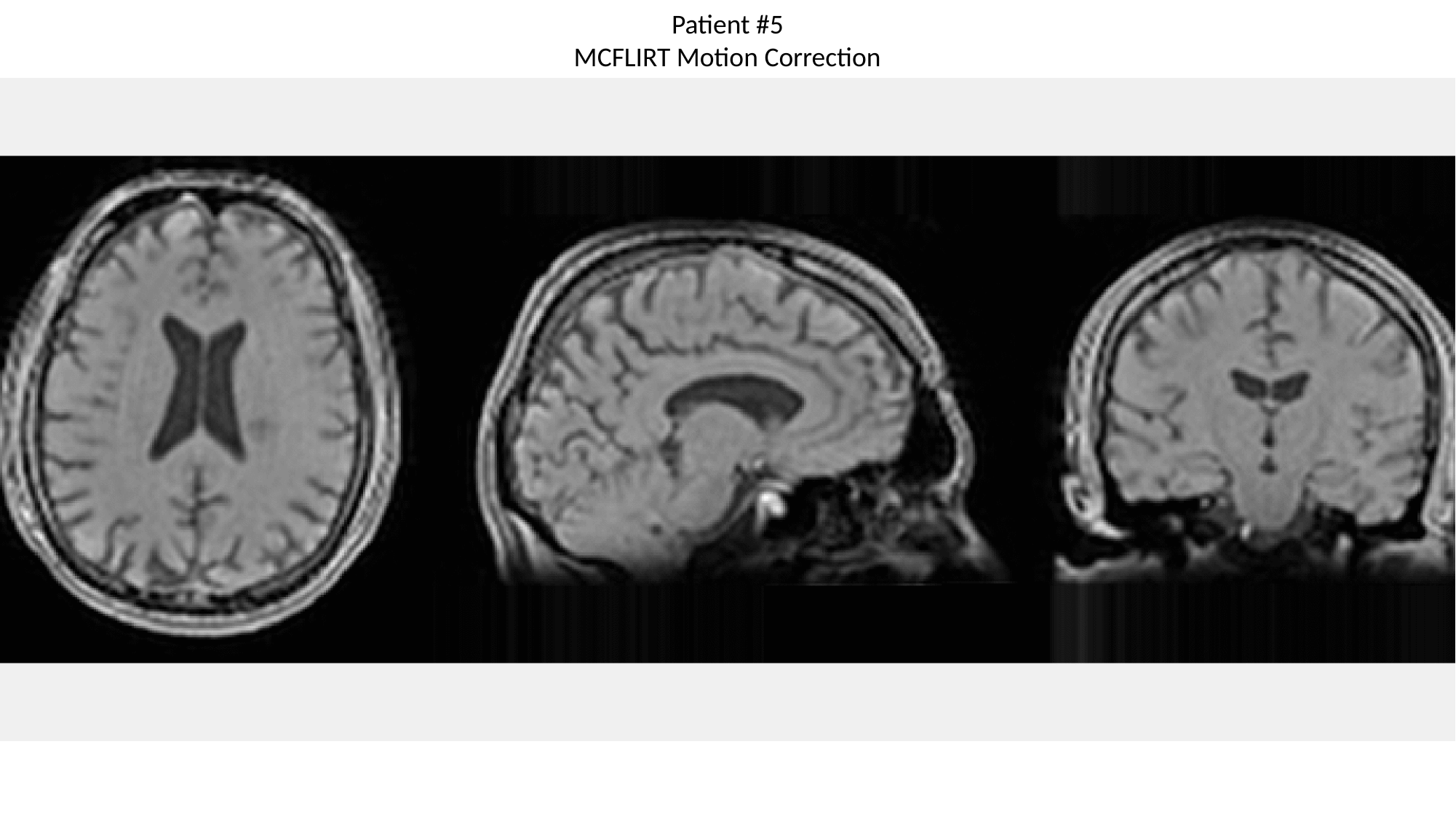

Patient #5
MCFLIRT Motion Correction

### Slide 15
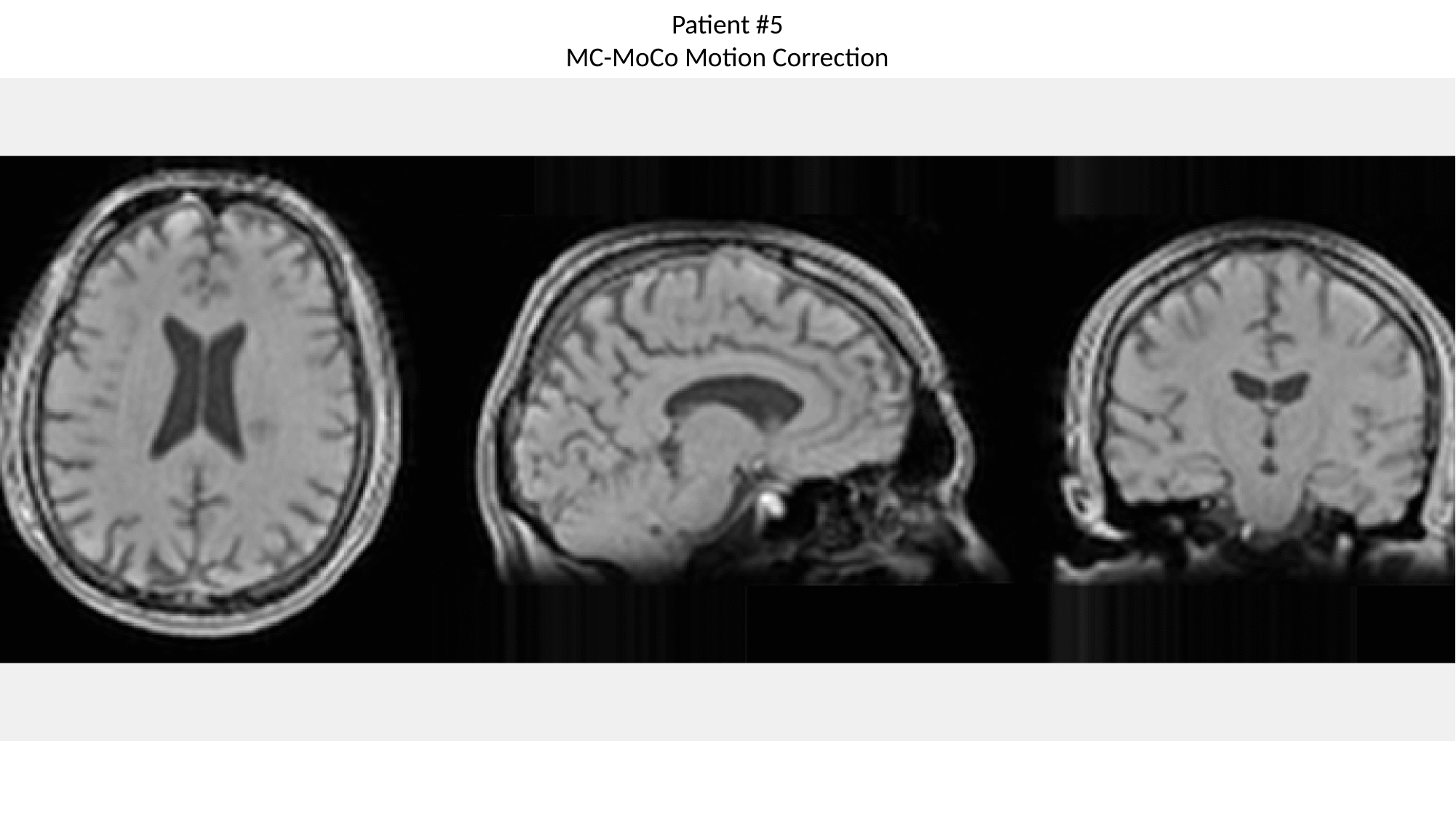

Patient #5
MC-MoCo Motion Correction

### Slide 16
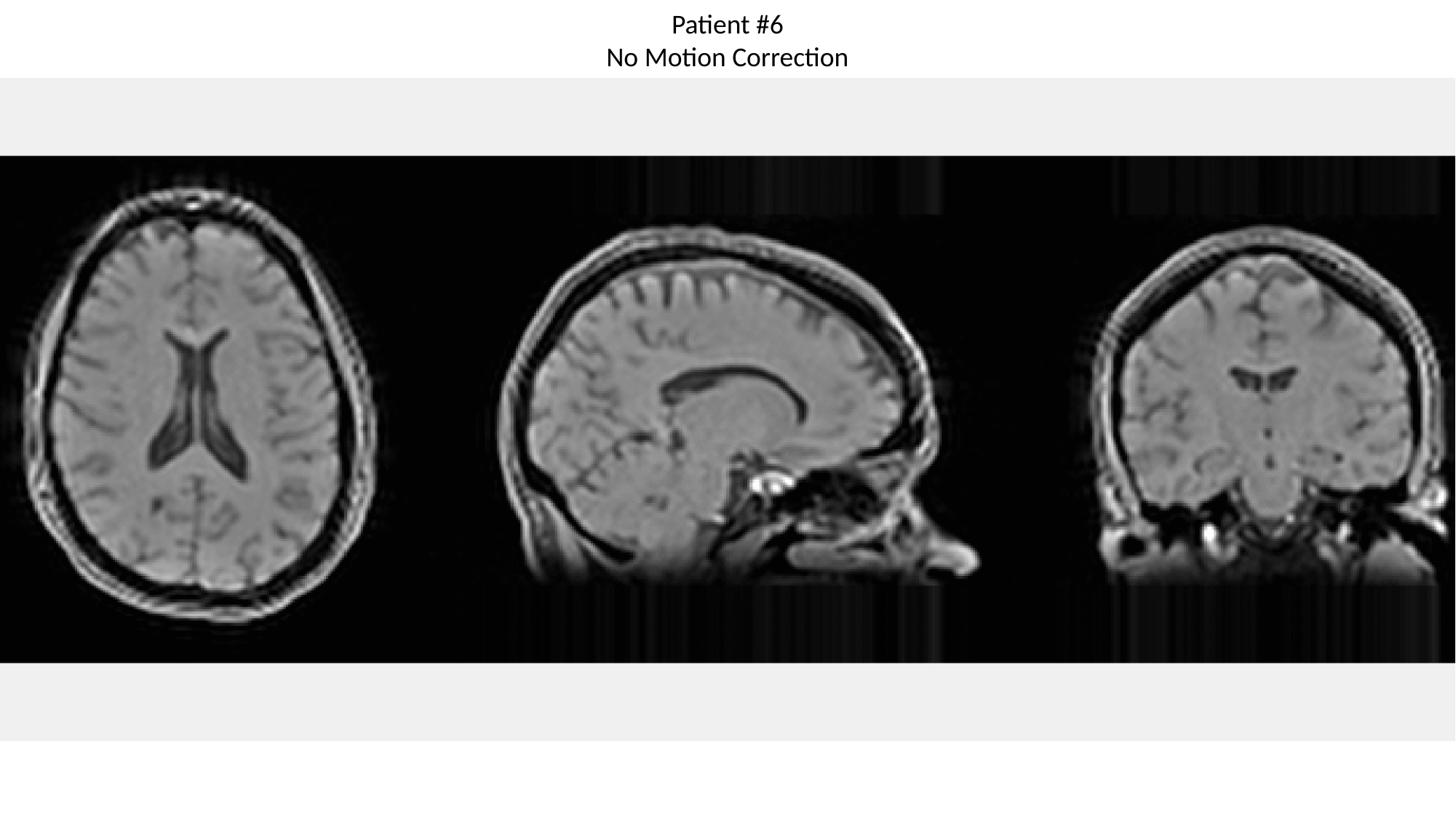

Patient #6
No Motion Correction

### Slide 17
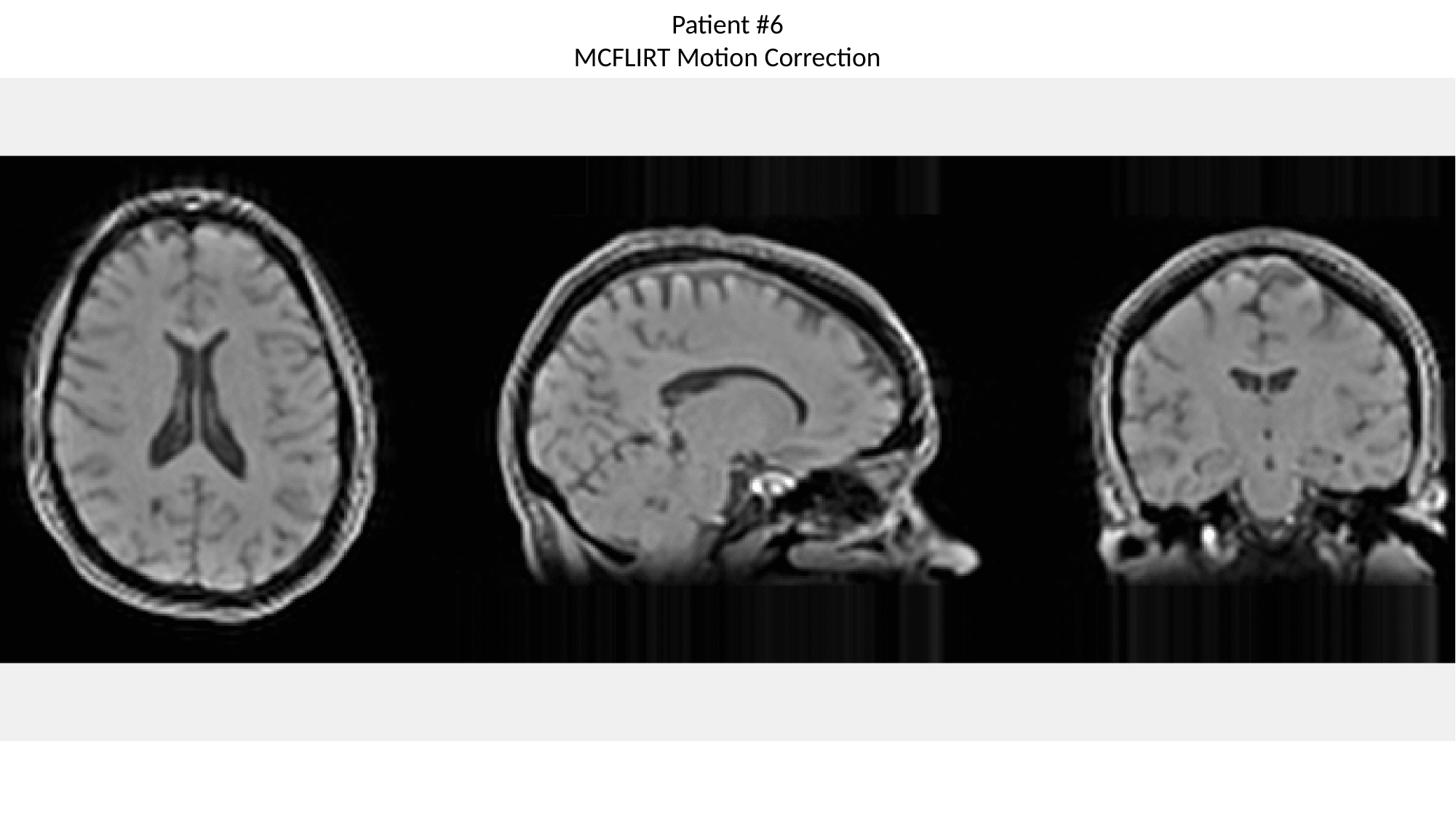

Patient #6
MCFLIRT Motion Correction

### Slide 18
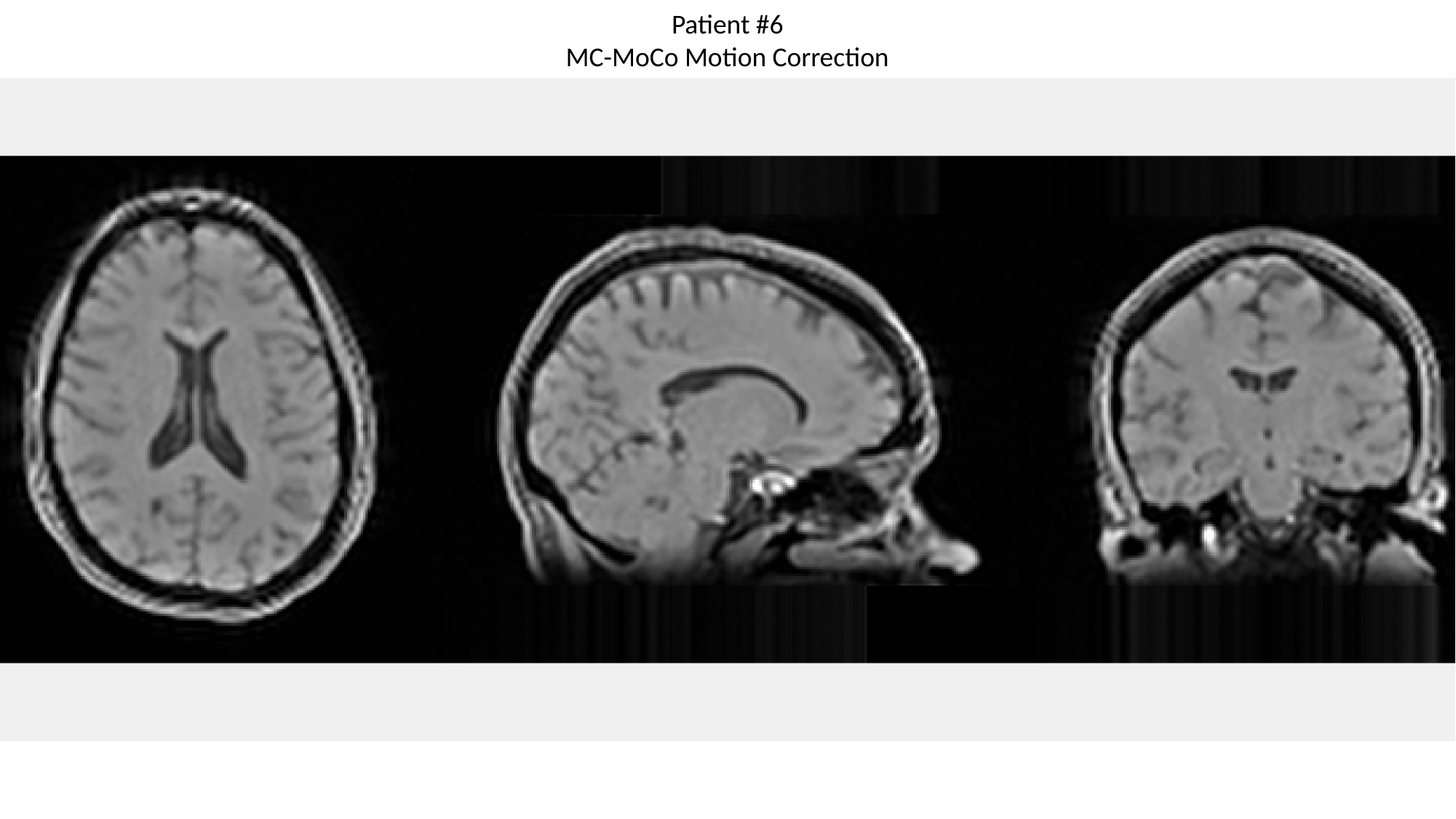

Patient #6
MC-MoCo Motion Correction

### Slide 19
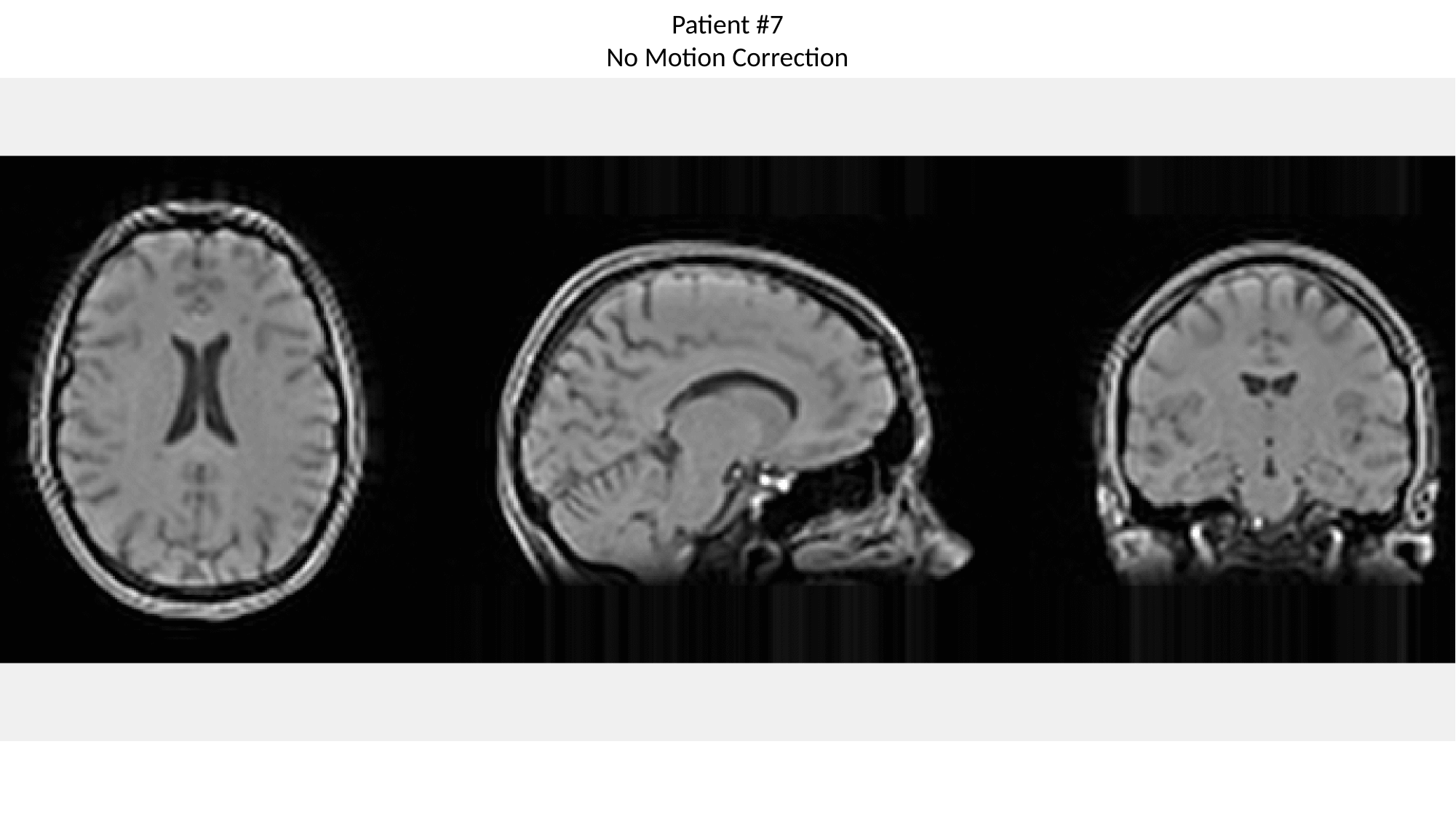

Patient #7
No Motion Correction

### Slide 20
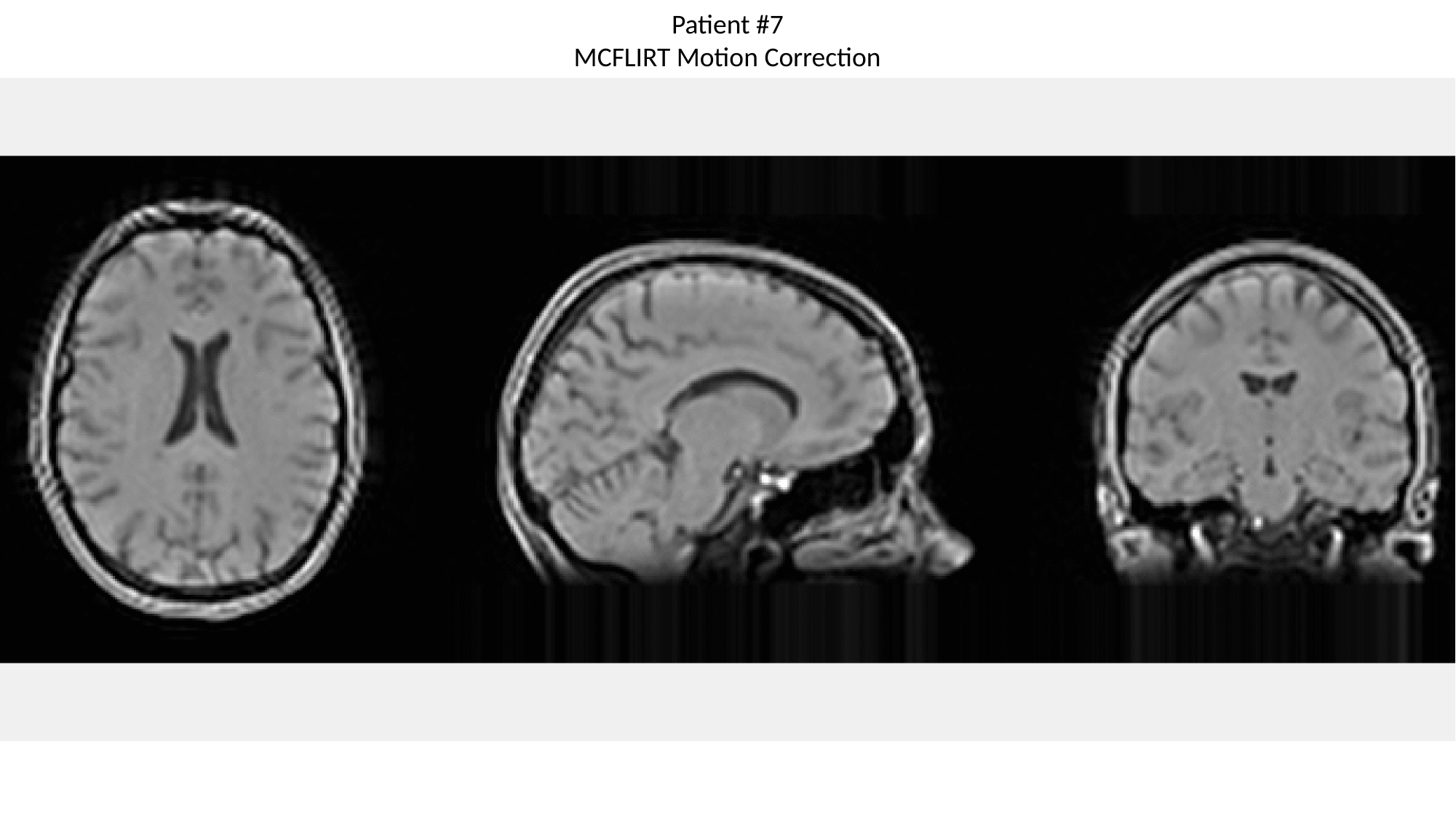

Patient #7
MCFLIRT Motion Correction

### Slide 21
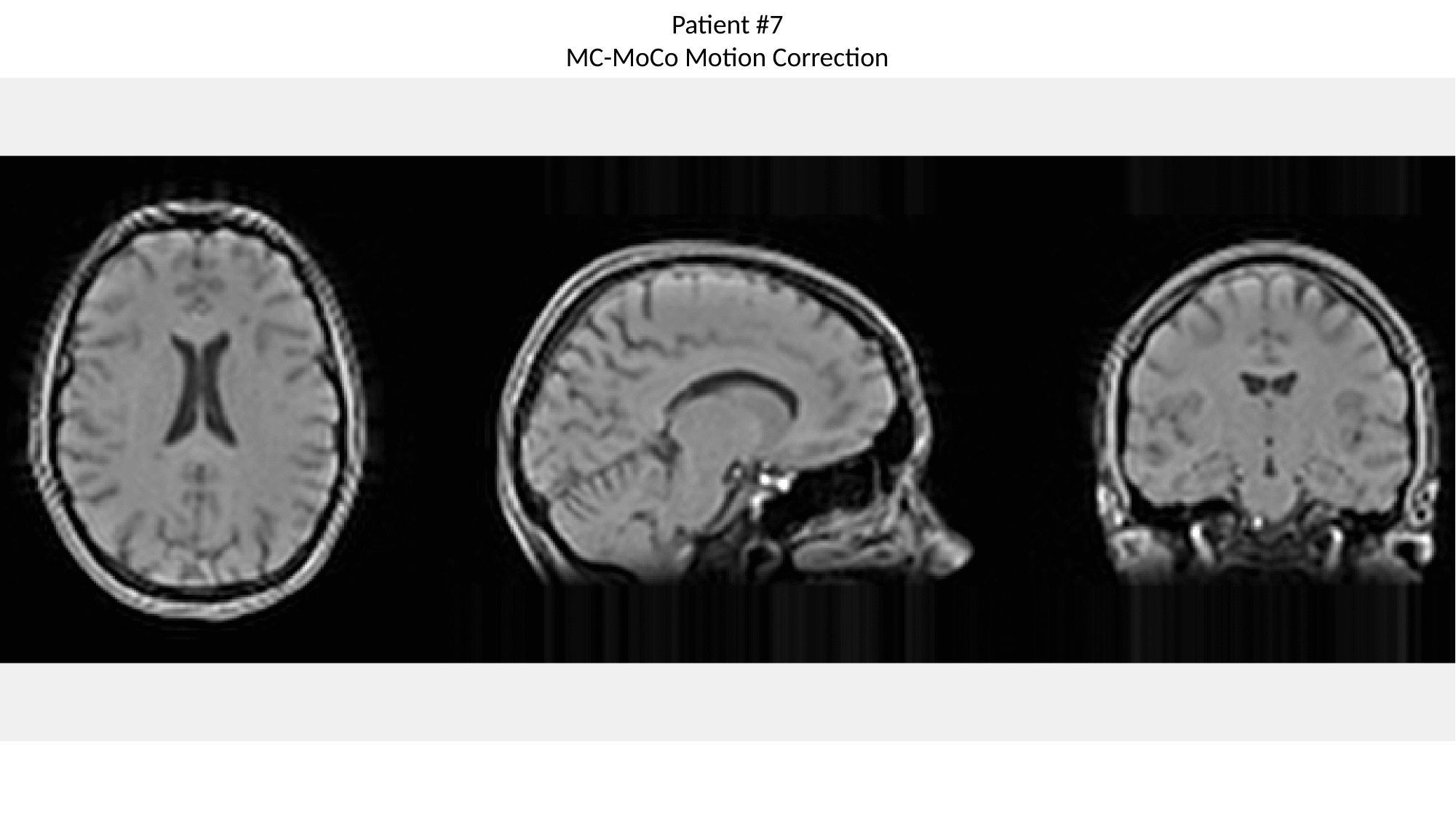

Patient #7
MC-MoCo Motion Correction

### Slide 22
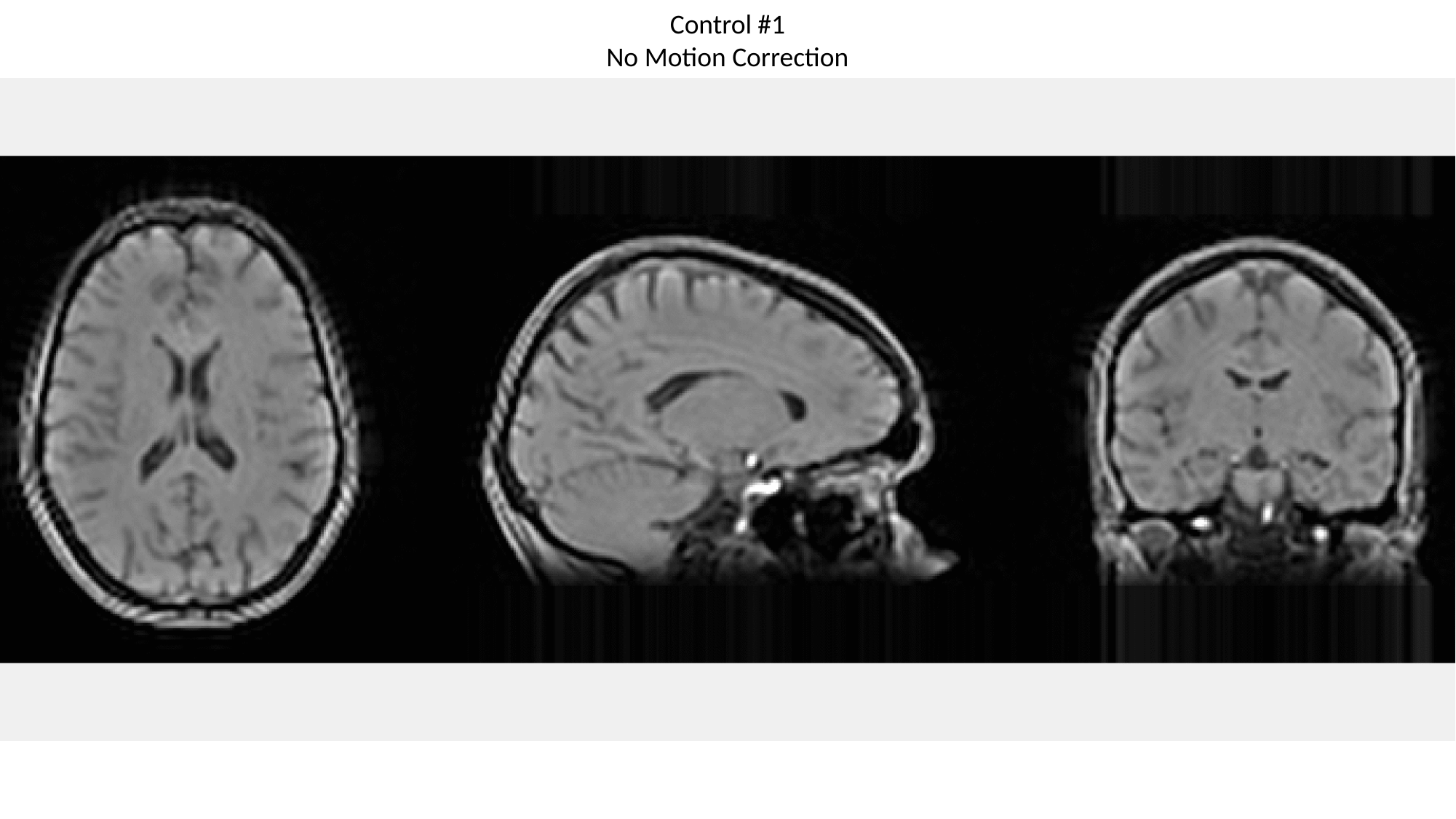

Control #1
No Motion Correction

### Slide 23
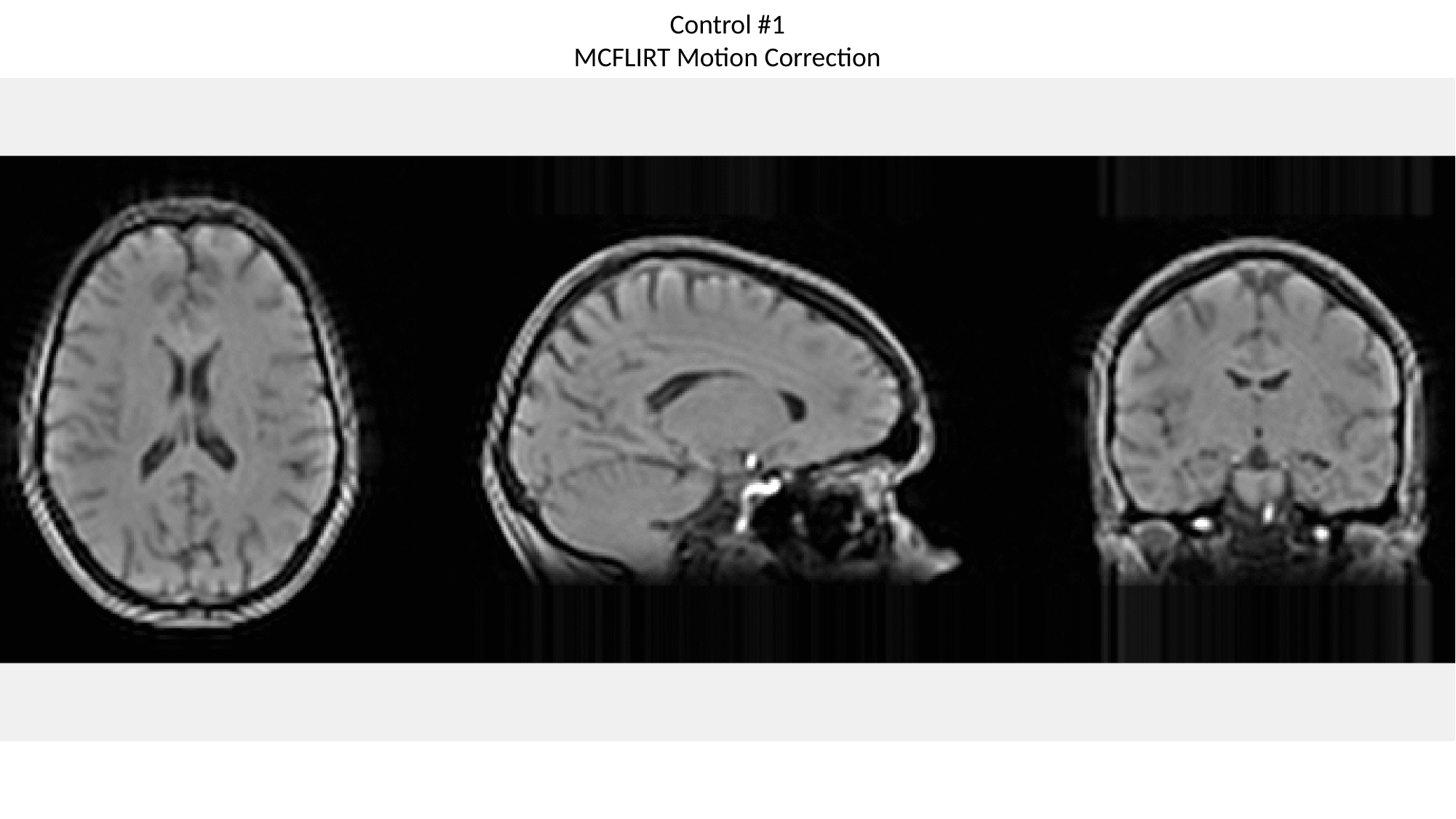

Control #1
MCFLIRT Motion Correction

### Slide 24
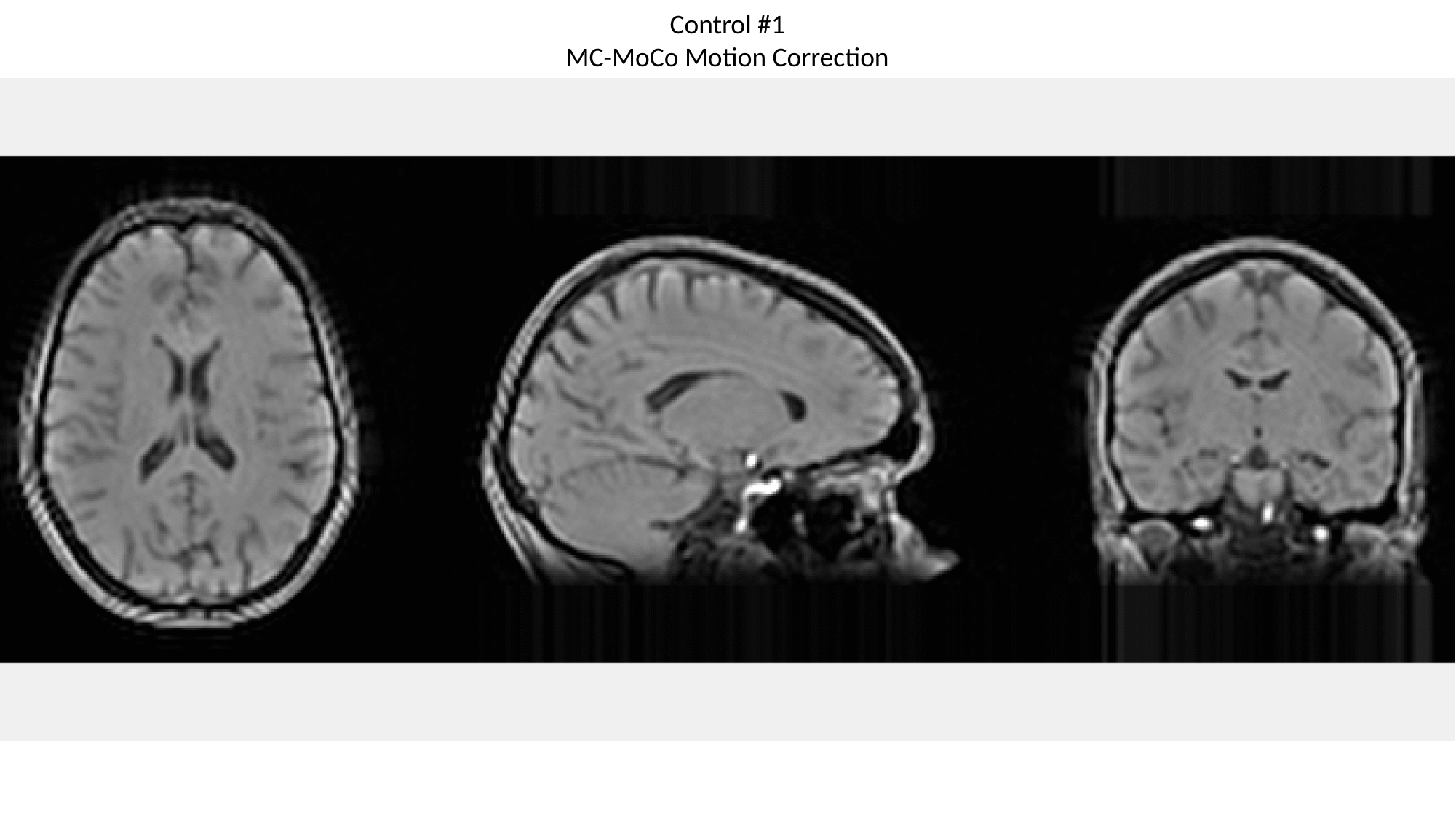

Control #1
MC-MoCo Motion Correction

### Slide 25
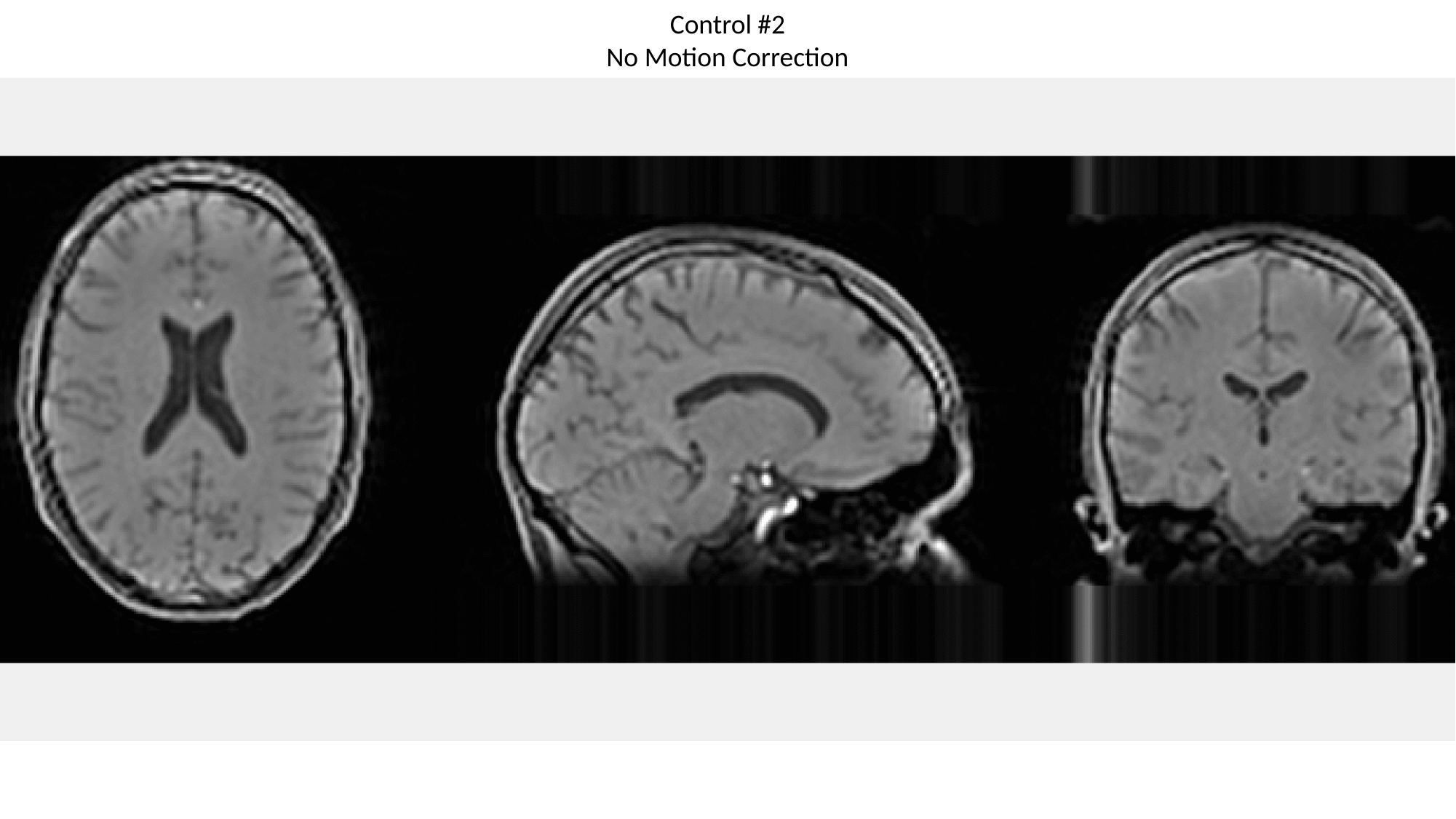

Control #2
No Motion Correction

### Slide 26
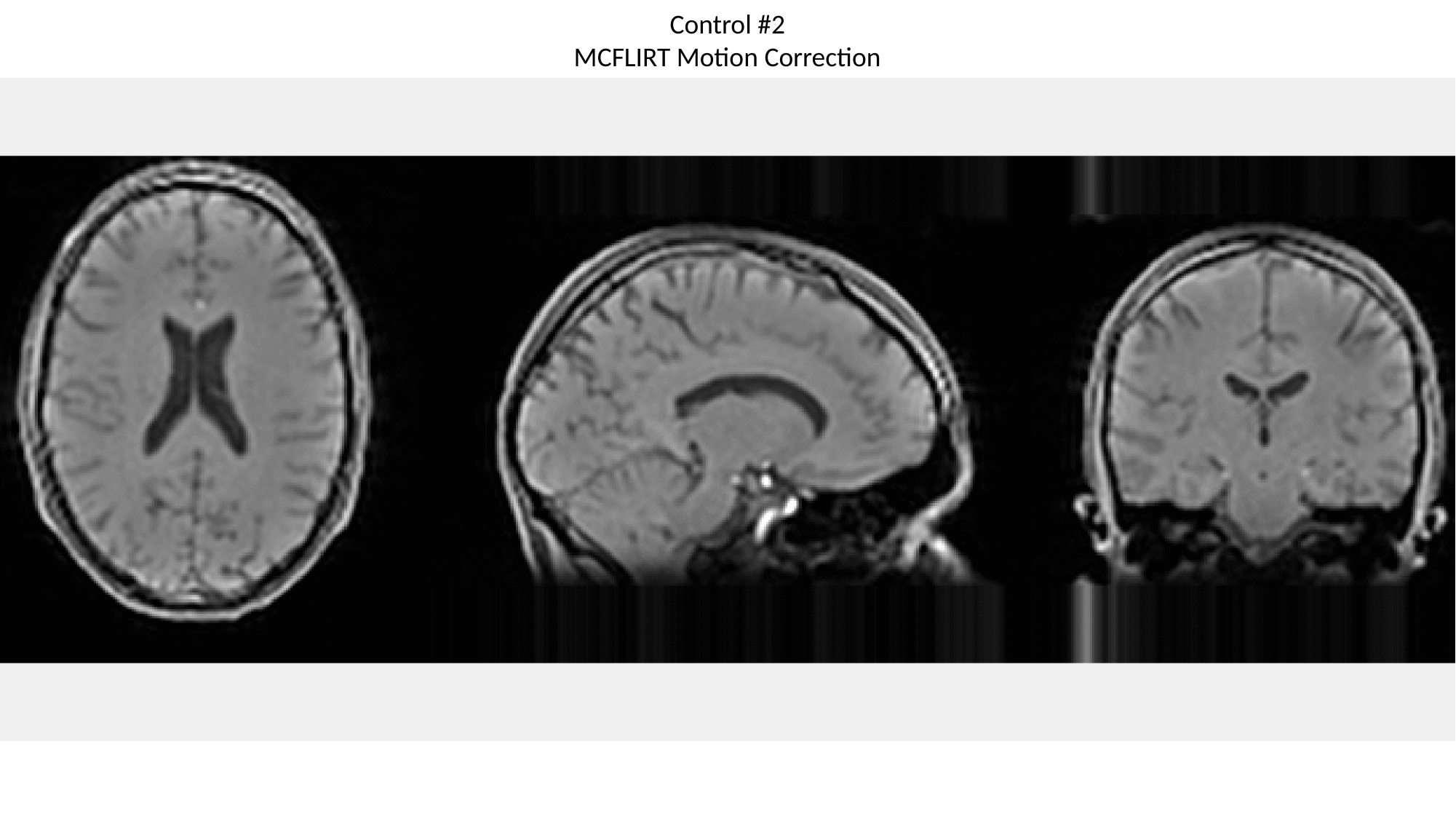

Control #2
MCFLIRT Motion Correction

### Slide 27
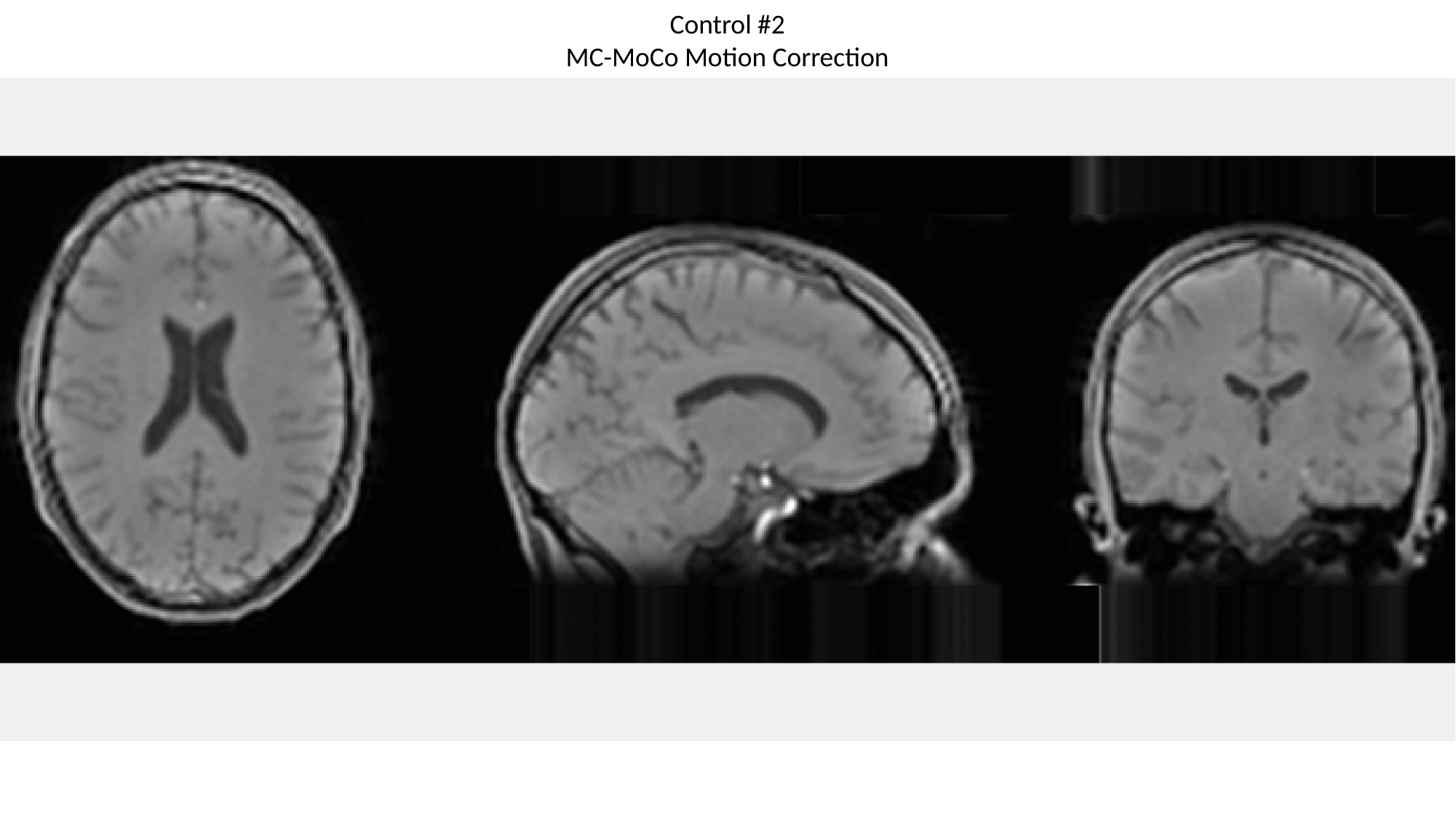

Control #2
MC-MoCo Motion Correction

### Slide 28
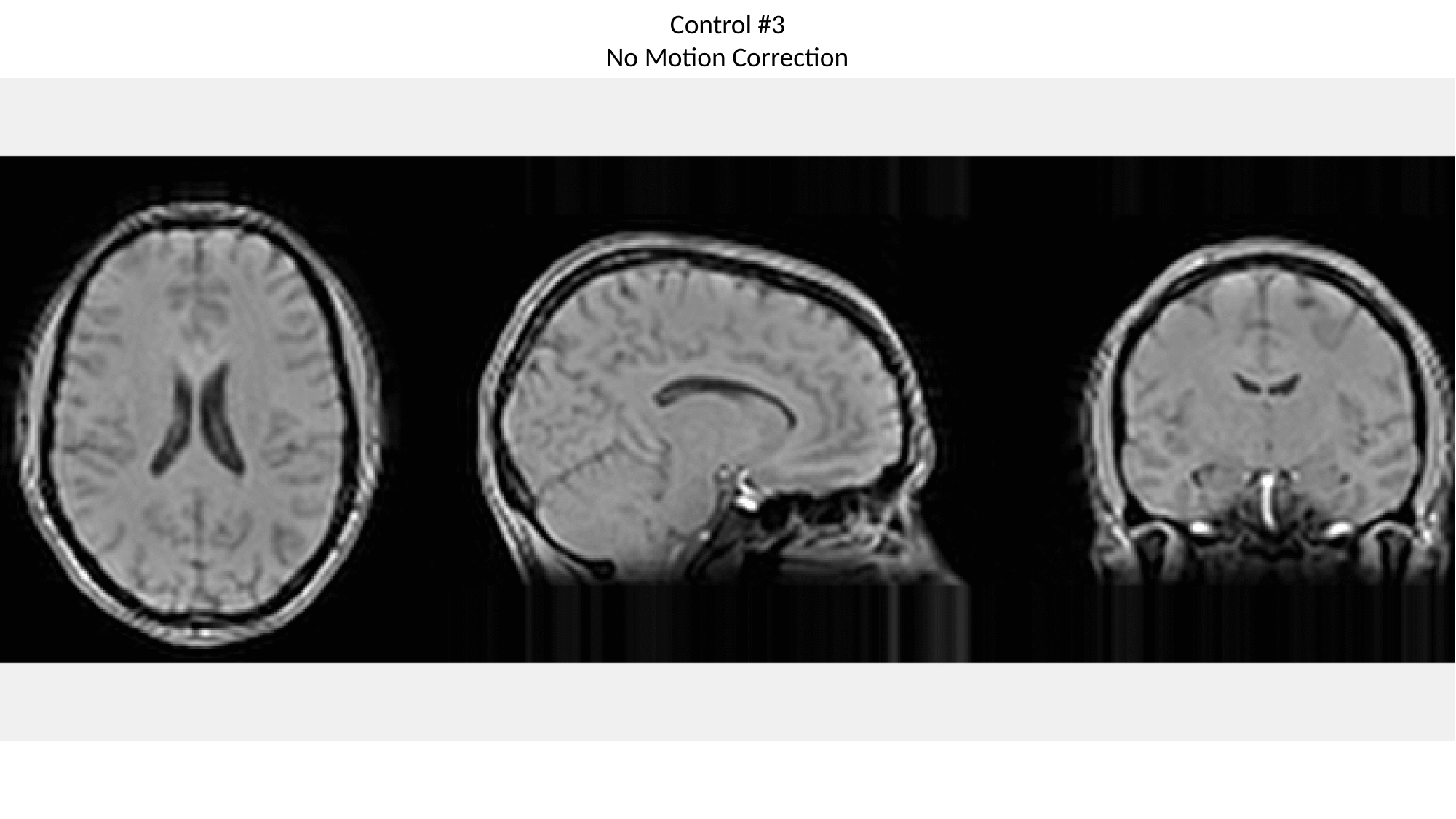

Control #3
No Motion Correction

### Slide 29
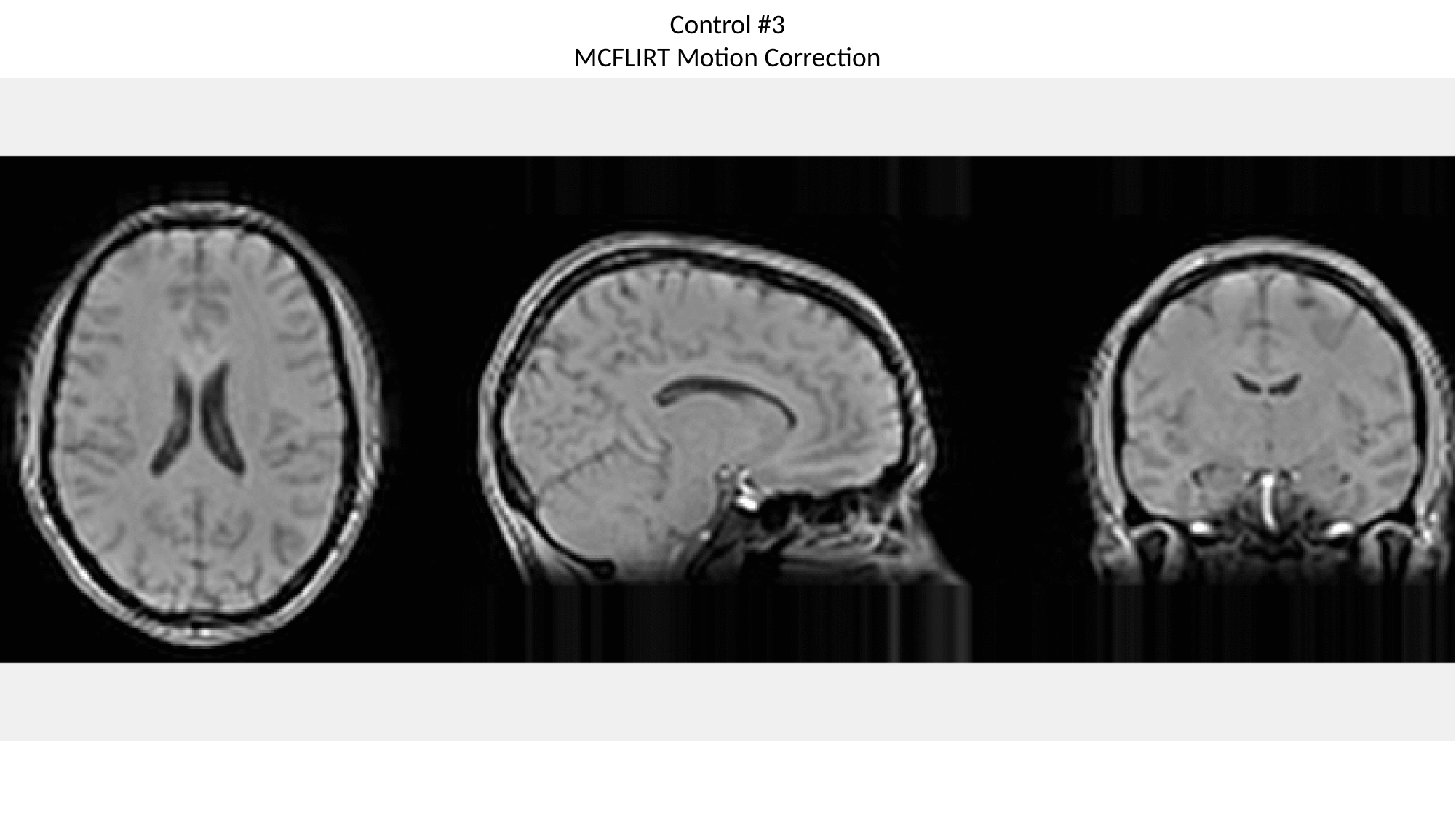

Control #3
MCFLIRT Motion Correction

### Slide 30
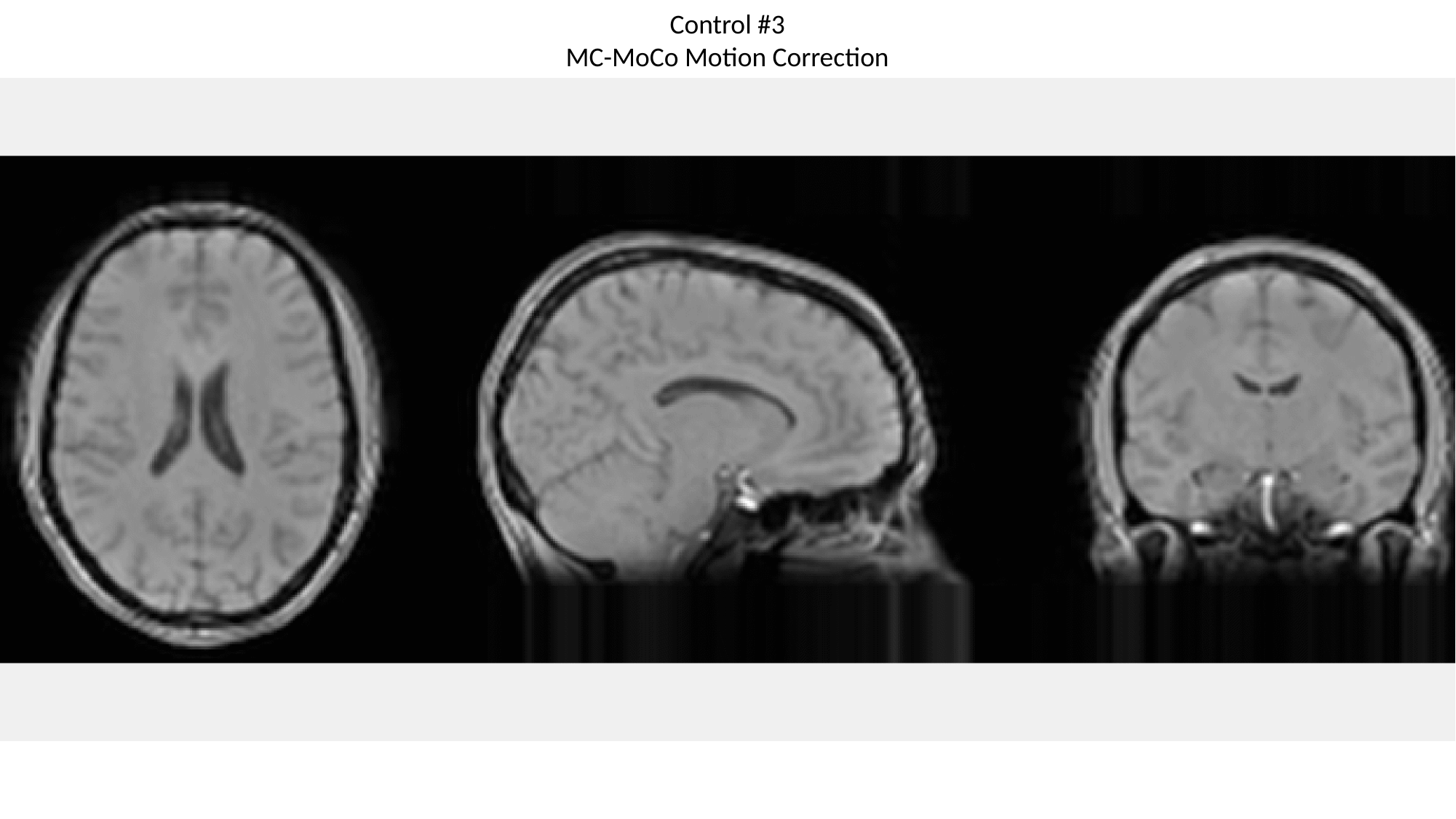

Control #3
MC-MoCo Motion Correction

### Slide 31
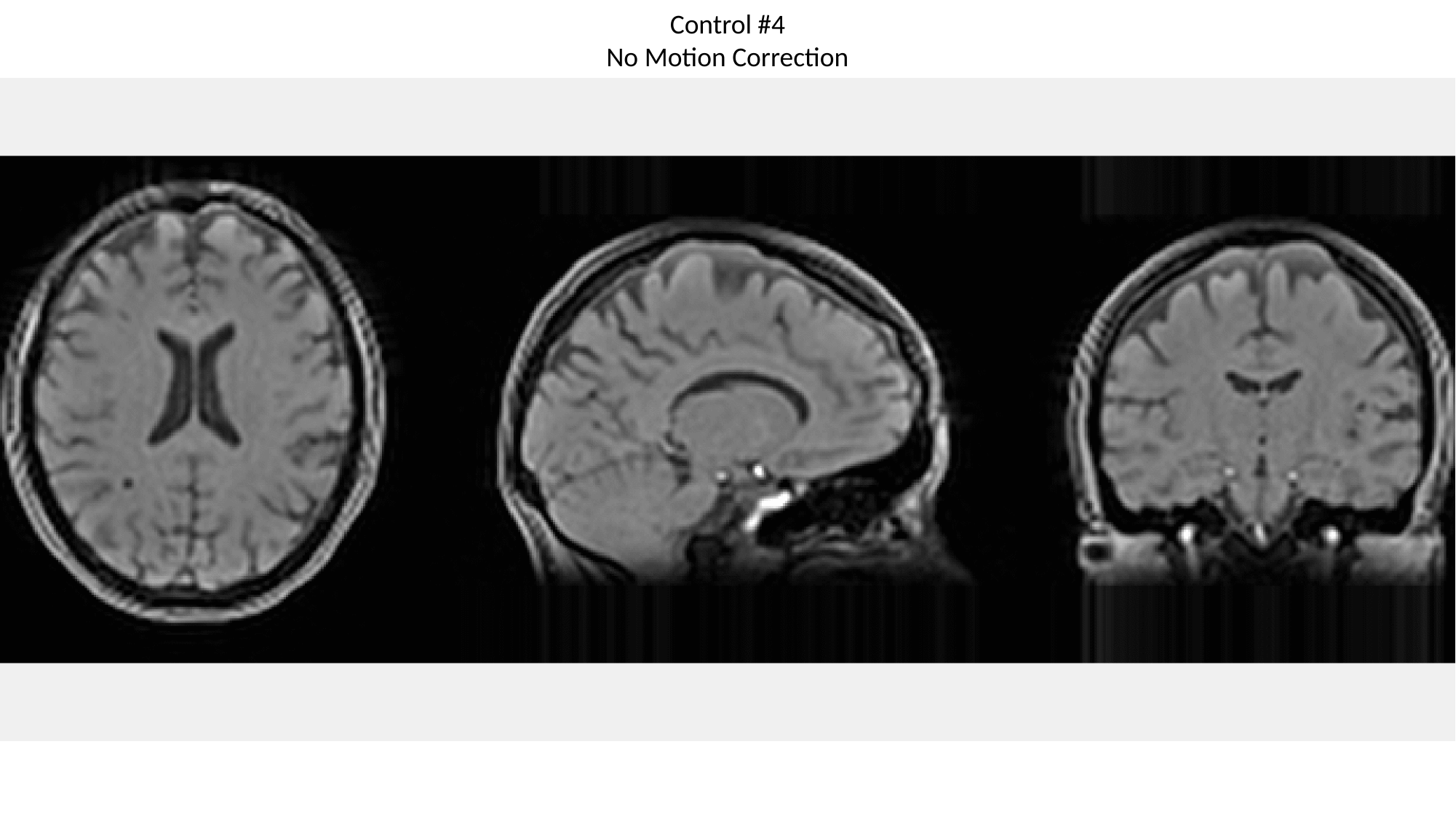

Control #4
No Motion Correction

### Slide 32
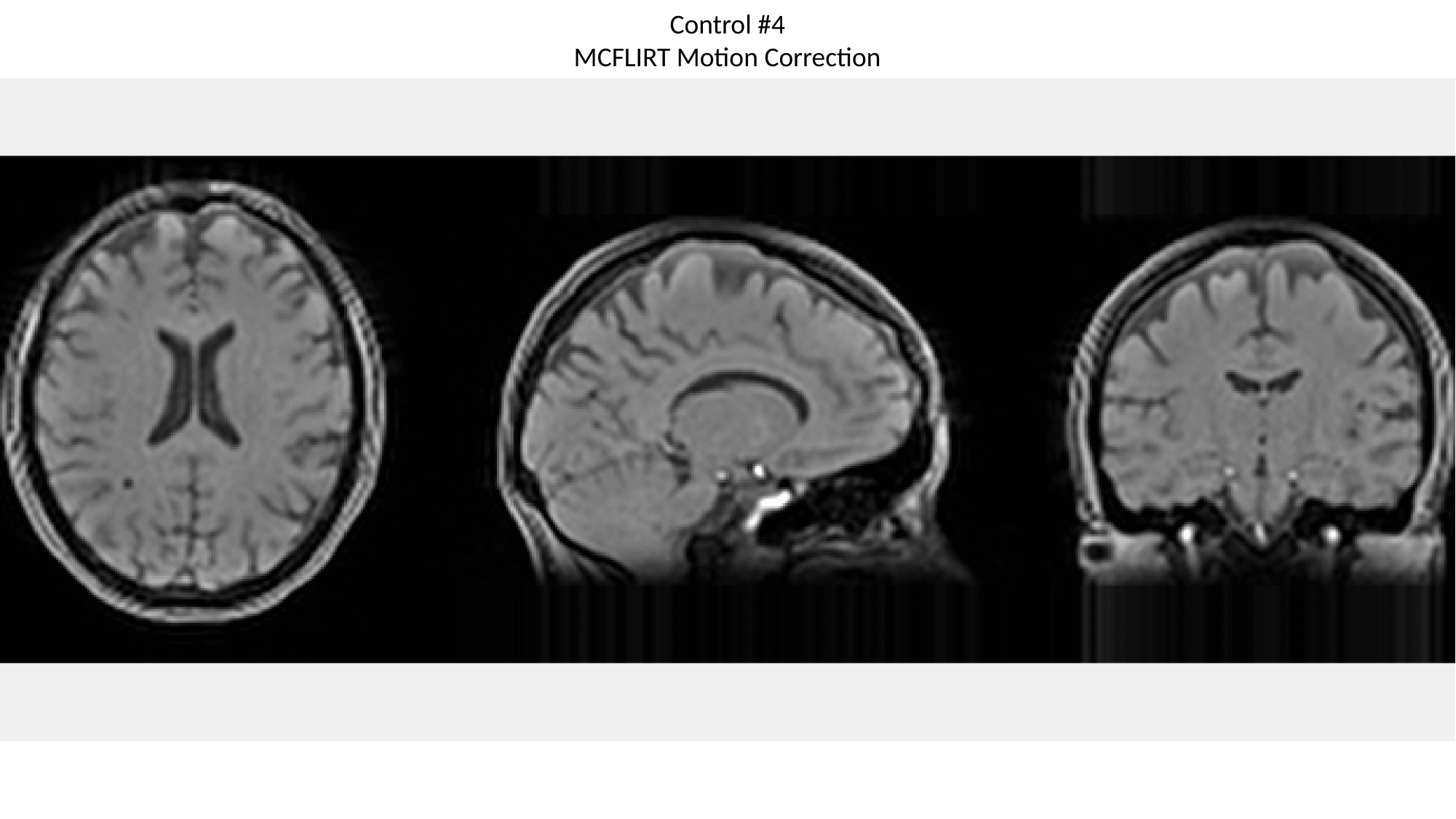

Control #4
MCFLIRT Motion Correction

### Slide 33
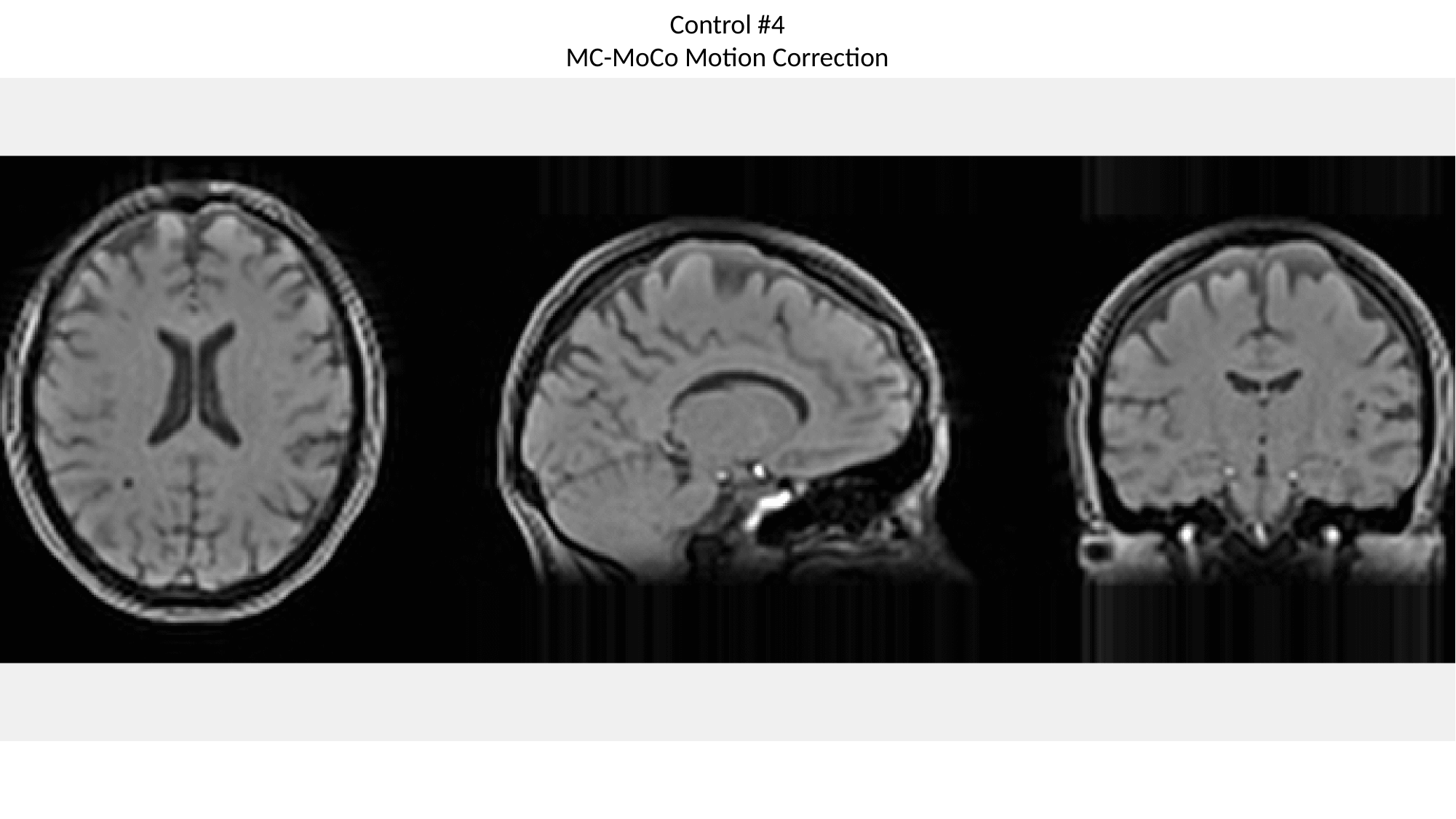

Control #4
MC-MoCo Motion Correction

### Slide 34
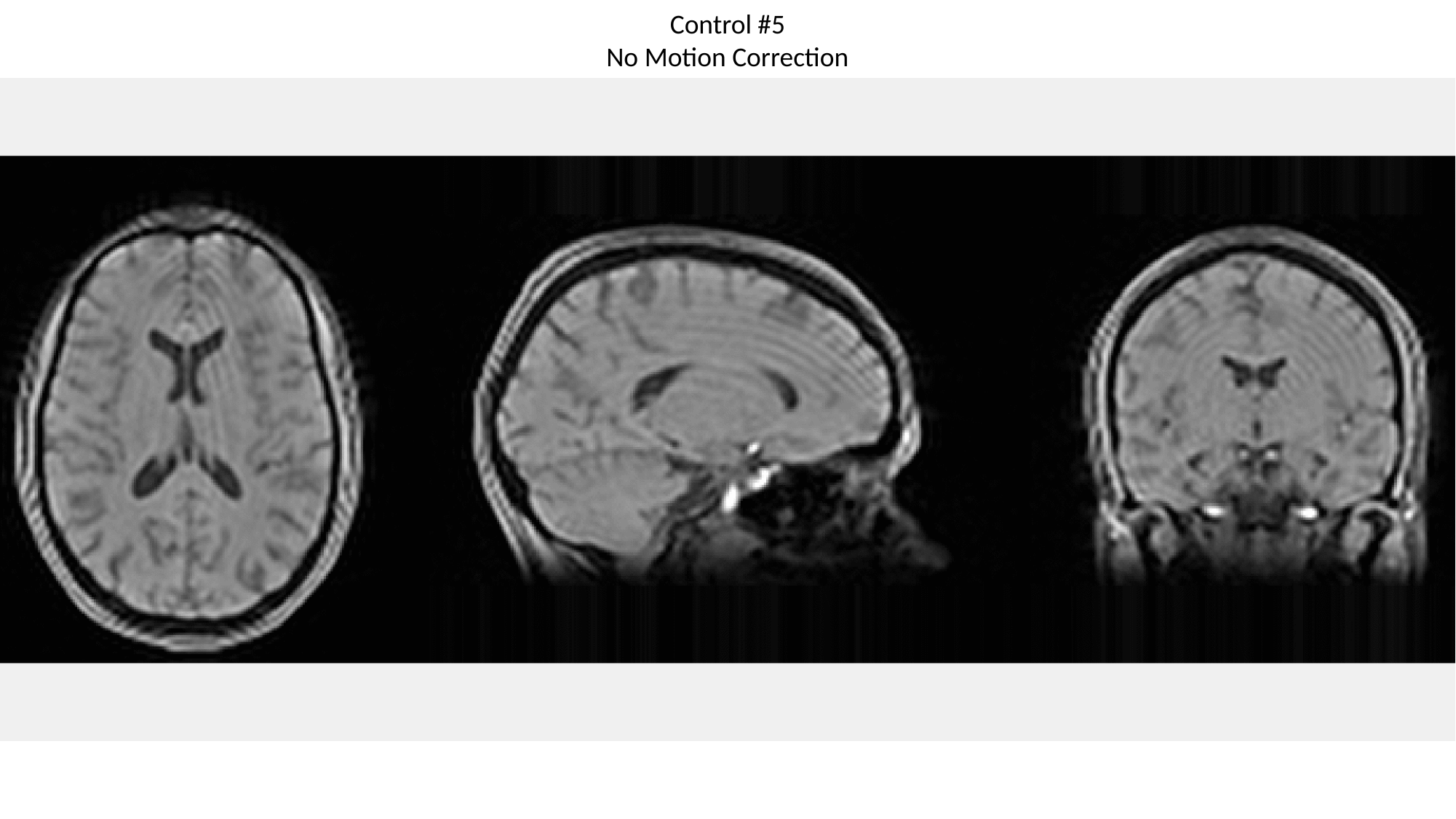

Control #5
No Motion Correction

### Slide 35
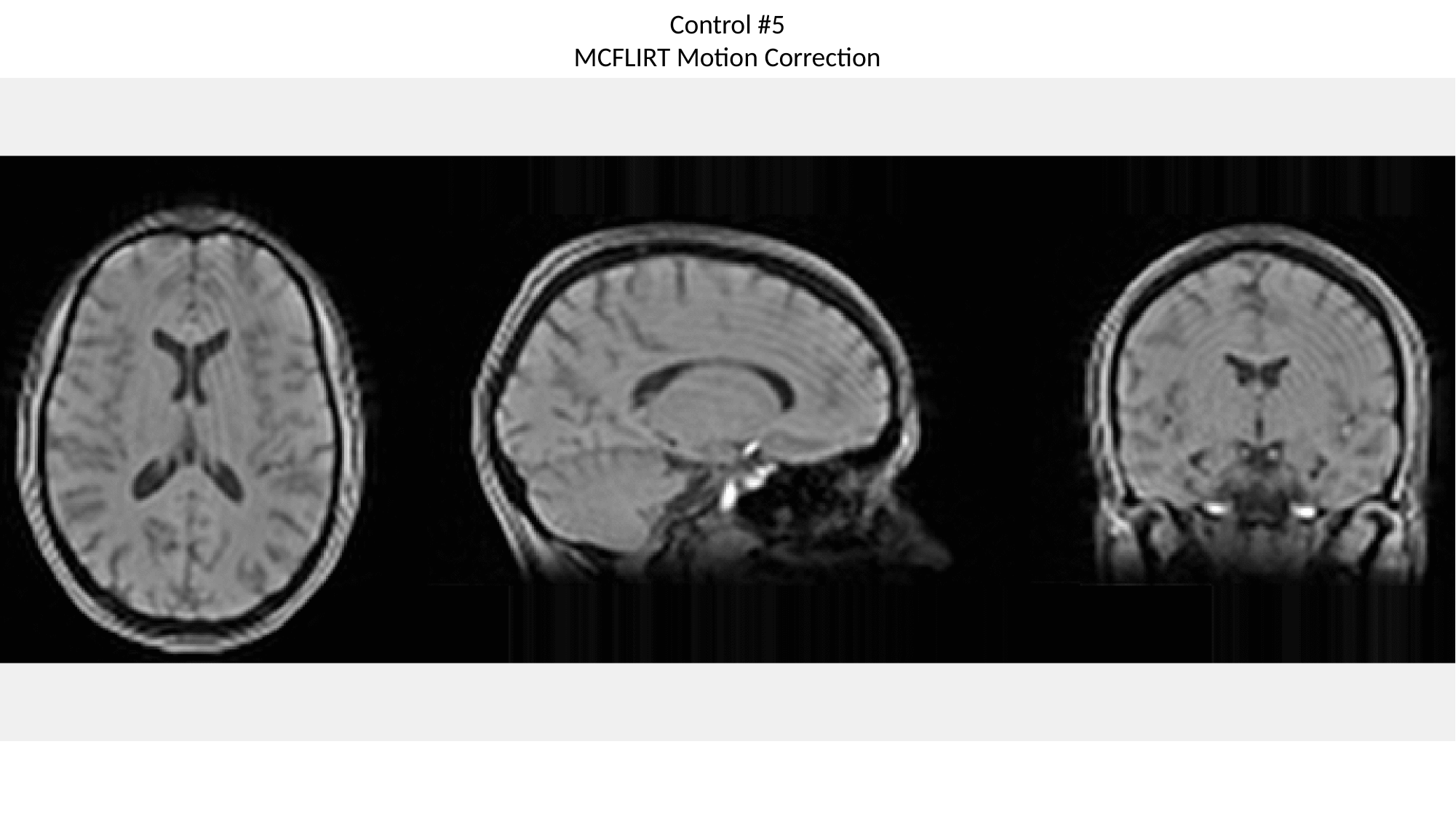

Control #5
MCFLIRT Motion Correction

### Slide 36
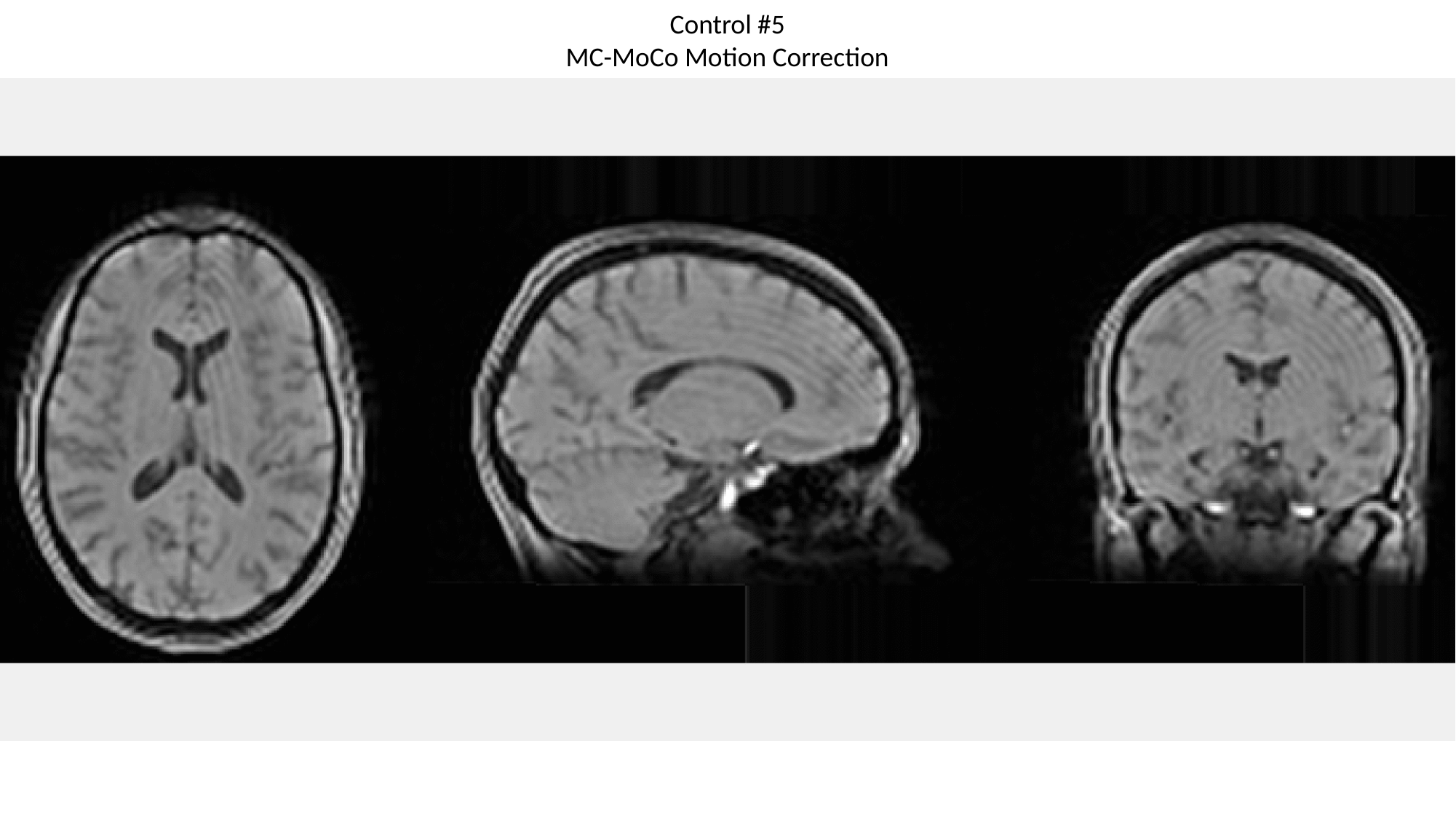

Control #5
MC-MoCo Motion Correction

### Slide 37
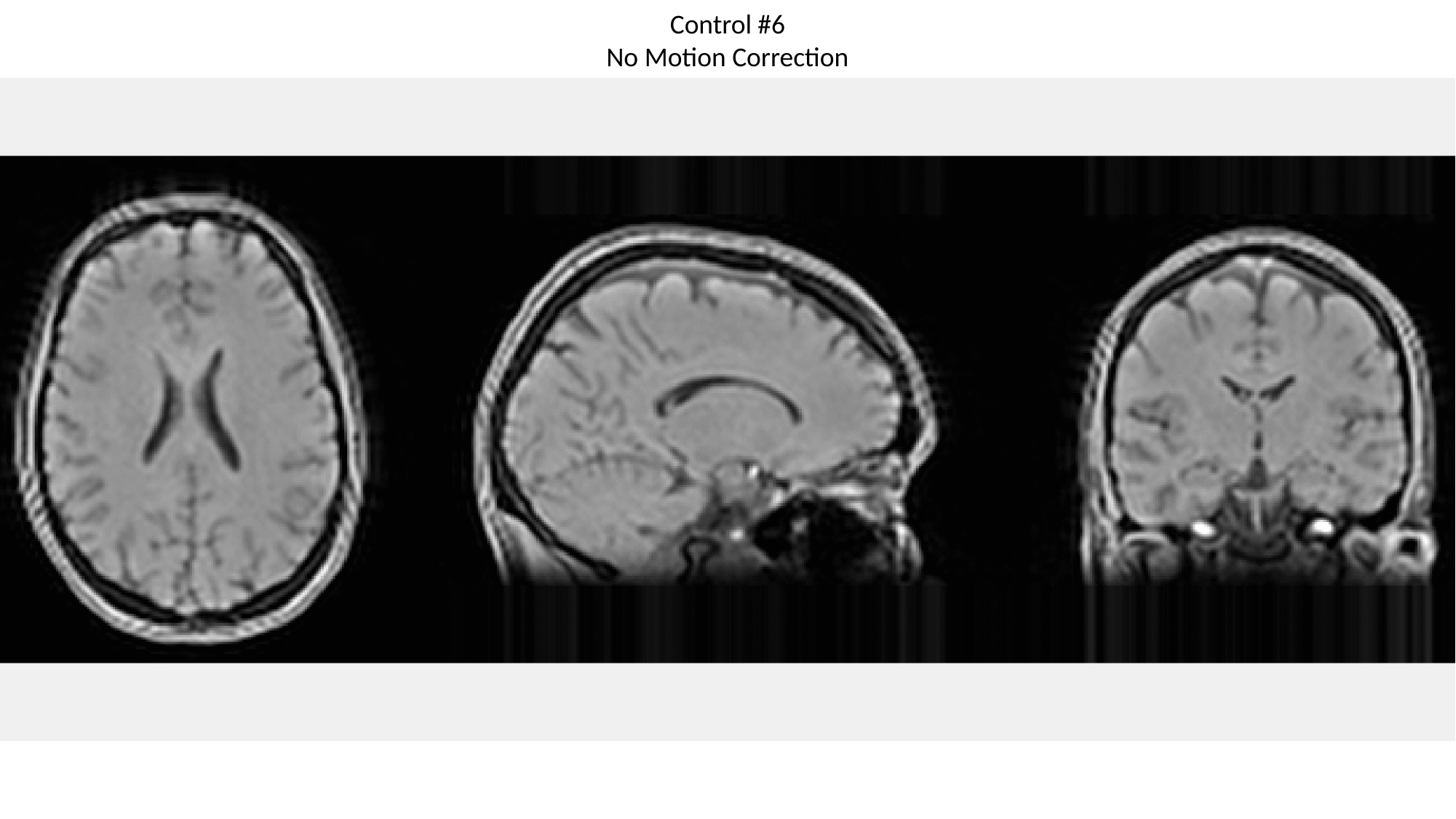

Control #6
No Motion Correction

### Slide 38
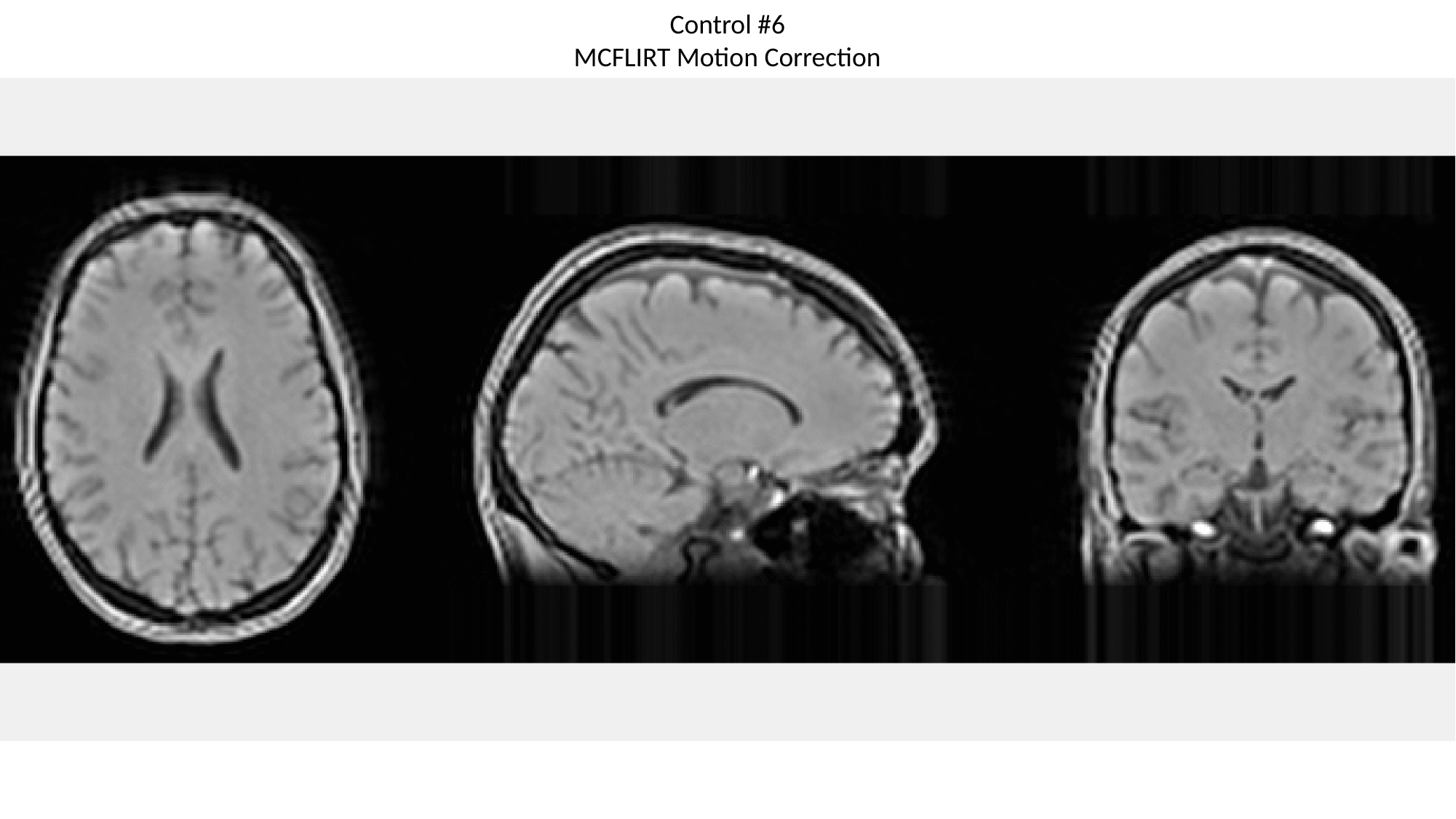

Control #6
MCFLIRT Motion Correction

### Slide 39
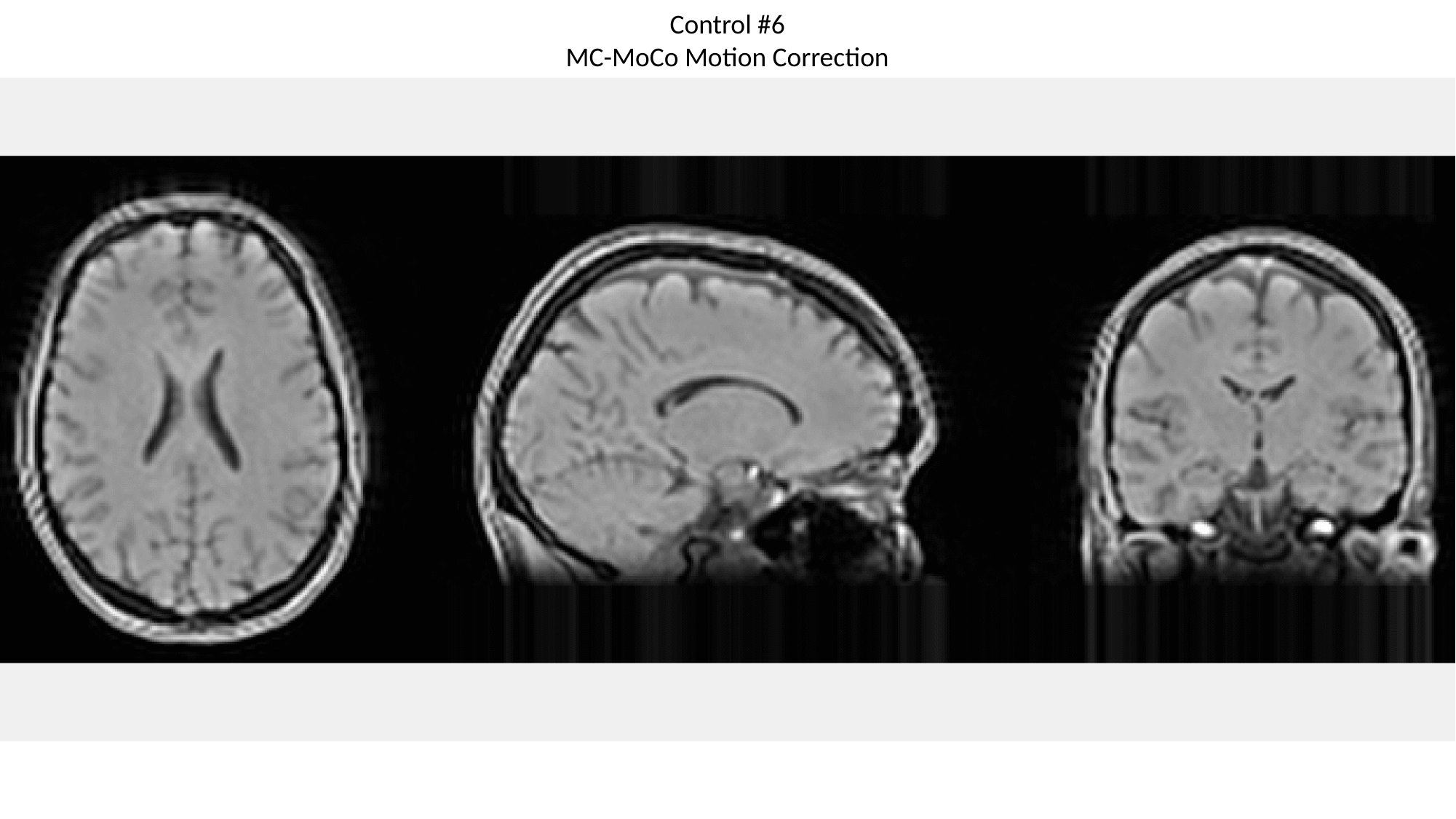

Control #6
MC-MoCo Motion Correction
